# Evolutionary conserved aspects of animal nutrient uptake and transport in sea anemone vitellogenesis

**DOI:** 10.1101/2022.01.24.477498

**Authors:** Marion Lebouvier, Paula Miramón-Puértolas, Patrick R. H. Steinmetz

**Affiliations:** Sars International Centre for Marine Molecular Biology, University of Bergen, Thormøhlensgate 55, 5006 Bergen, Norway

## Abstract

Vitellogenesis, the accumulation of egg yolk, relies on the transport of dietary nutrients from the gut to the ovary through the circulatory system in many bilaterians (e.g. vertebrates, arthropods). How these dietary nutrients and yolk precursors are absorbed and transported in cnidarians (e.g. corals, sea anemones, jellyfish), which are bi-layered and lack a circulatory system, is however only poorly understood. Here, we studied the tissues and molecules that facilitate the uptake and transport of dietary nutrients, especially lipids, towards the oocytes in the sea anemone *Nematostella vectensis* to better understand the evolution of systemic nutrient transport in animals. We identified the somatic gonad epithelium as one of several gastrodermal tissues specialized in phagocytosis, micropinocytosis and intracellular digestion. We showed more specifically that dietary fatty acids are absorbed by the ApolipoproteinB- and Vitellogenin-expressing somatic gonad epithelium. Their subsequent, rapid transport into the extracellular matrix (ECM) and endocytosis into oocytes is likely mediated by an evolutionary conserved Vitellogenin (Vtg)-Very Low-Density Lipoprotein Receptor (VLDLR) ligand/receptor pair. We propose that ECM-based, Vtg/VLDLR-mediated lipoprotein transport during vitellogenesis predates the cnidarian-bilaterian split and provided a mechanistic basis to evolve sophisticated circulatory systems in bilaterians.

## Introduction

Dietary nutrient transport is essential for animal energy homeostasis and gametogenesis^1^. In bilaterians, ApolipoproteinB (ApoB; termed Apolipophorins in insects) and Vtg, two proteins of the Large Lipid Transfer Protein (LLTP) superfamily, play a key conserved role during lipid transport^2,3^. ApoB mediates systemic lipid transport between the gut epithelium, vitellogenic organs and other peripheral tissues^4–6^. In contrast, Vtg specifically mediates lipid transport from vitellogenic organs into oocyte yolk^7–9^. Vtg expression is therefore mostly female-specific and primarily occurs in extra-ovarian tissues (e.g. vertebrate liver, insect fat body)^8,10^. In many bilaterians, Vtg and ApoB lipoproteins travel via the hemocoel or blood vascular system^11^ and are endocytosed into oocytes or other target tissues (e.g. muscles) by conserved orthologs of the VLDL receptor family, which also includes arthropod Vitellogenin or Apolipophorin receptors^12–14^. Although Vtg orthologs are almost ubiquitously present among animal phyla^15–17^, systemic lipoprotein transport is poorly studied in many animals, especially in non-bilaterians lacking a circulatory system. The evolution of complex bilaterian nutrient transport systems, including vertebrate or insect circulatory systems, remains therefore speculative^18–20^.

Adult *Nematostella* polyps exhibit a typical bi-layered cnidarian body plan, consisting of the inner gastrodermis and outer epidermis epithelia. Food uptake occurs through the mouth into the sack-like gastric cavity, which is therefore commonly assumed to constitute the main nutrient distribution system in cnidarians^21^. Gastrodermal folds (mesenteries) reach into the gastric cavity of anthozoans (sea anemones, corals, sea pens) - including *Nematostella*. They are likely an ancestral feature of cnidarians^22,23^ and are subdivided into functionally distinct regions (Fig. 1a)^24–26^. Within the oral half of the *Nematostella* body column, oocytes or spermaries are embedded in extracellular matrix, the mesoglea, between two layers of somatic gonad epithelia, collectively termed ‘gonad’^26,27^. The ECM localization of gametes is common among anthozoans (sea anemones, corals, sea pens), scyphozoans (‘true’ jellies), cubozoans (cube jellies) and hydrozoan medusae^21,28^ and thus likely ancestral in cnidarians. Cnidarian germ cells are therefore most often not in direct contact with food but rely on nutrient transport through the extracellular matrix.

**Figure 1:**
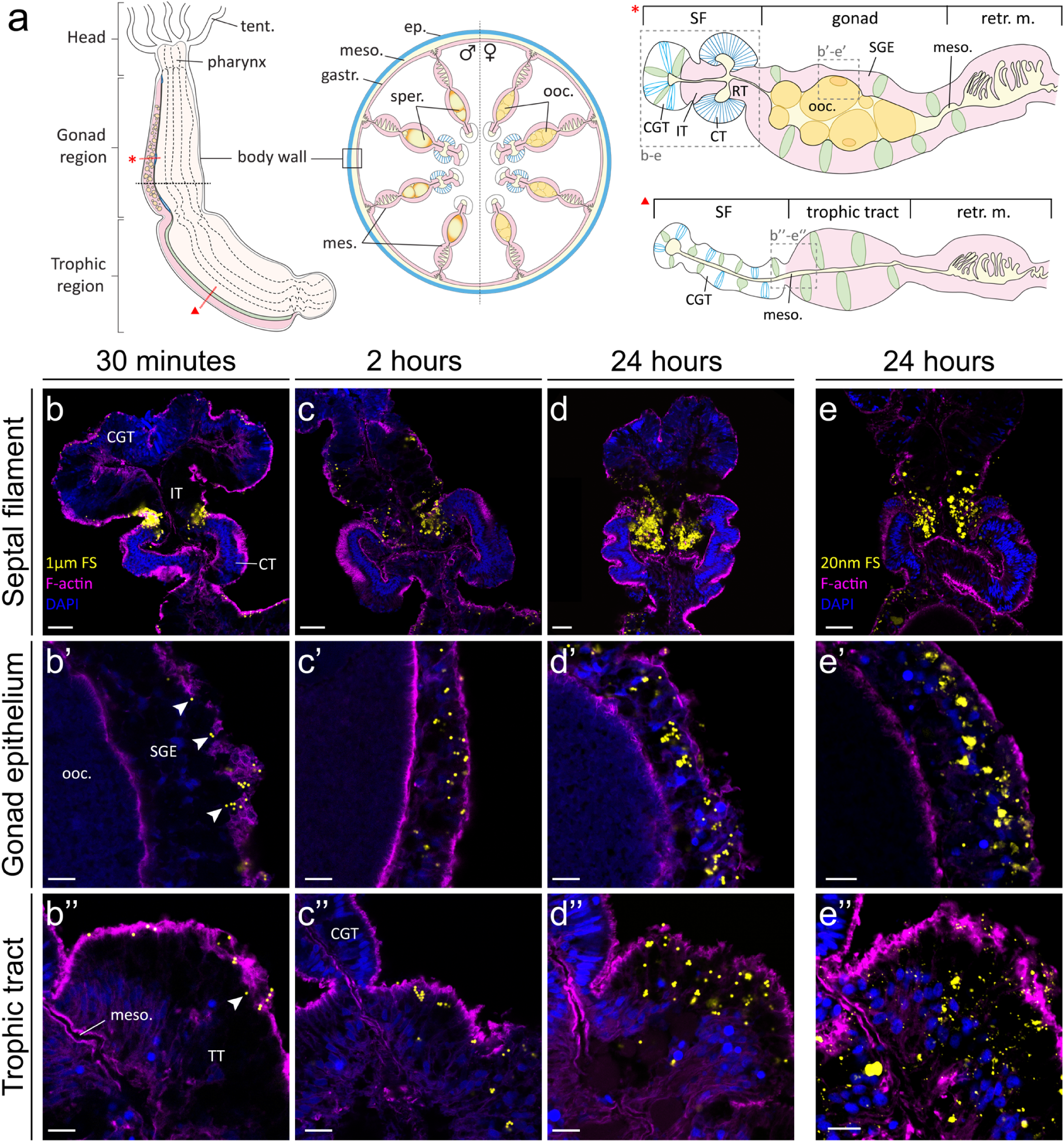
Regionally specific uptake of 1μm and 20nm Fluospheres (FS) by mesenterial epithelium. (a) Schematic representation of *Nematostella* morphology. (b-e”) Specific uptake of 1μm (b-d”) or 20nm (e-e”) Fluospheres (yellow) by specific parts of the mesenteries. 1μm FS are progressively internalized into cells of the intermediate tract (b, c, d, e), gonad epithelium (b’, c’, d’, e’) and trophic tract (b”, c”, d”, e”) after 30 minutes (b-b’’), 2 hours (c-c’’) and 24 hours (d-d’’) of incubation. A similar distribution of particle uptake is seen when incubating with 20nm FS (e-e’’). Distal end of mesentery oriented towards top (b-e”). All data figures represent single confocal plane images of adult *N. vectensis* mesenterial cross sections. Arrowheads in (b’, b’’): intracellular beads as judged by their relative location to cortical F-actin enrichment. Magenta: Phalloidin (F-actin); blue: DAPI. Scale bars: (b-e) 25μm, (e’-e’’) 10 μm. CGT: cnidoglandular tract, CT: ciliated tract, ep.: epidermis, gastr.: gastrodermis, IT: intermediate tract, mes.: mesentery, meso.: mesoglea, ooc.: oocyte, retr. m.: retractor muscle, SGE: somatic gonad epithelium, SF: septal filament, tent.: tentacle, TT: trophic tract.

How nutrients reach the oocyte from the gastric cavity and potentially other parts of the *Nematostella* body has been the focus of this study and is used as a paradigm for understanding the evolution of animal dietary lipid transport systems. The comparison with well-studied bilaterians, the phylogenetic sister group to cnidarians, allows the reconstruction of cellular and molecular aspects of the lipid transport system present in their last common ancestors.

## Results

### Specialized regions of the mesentery display high endocytic activities

The body regions where food particles or nutrients are absorbed are only poorly described in sea anemones^29–31^, and currently unknown in adult *Nematostella* polyps. We have used 1μm or 20nm BSA-coated fluorescent microspheres (Fluospheres) to probe for phagocytosis or micropinocytosis, respectively, in adult *Nematostella* polyps^32–34^. After incubation for 24 hours, we found 20nm beads but not 1μm beads enriched in the body wall epidermis suggesting that particles from the surrounding water are absorbed at high levels by micropinocytosis rather than phagocytosis in this region (Extended data fig. 1a, c). Both bead sizes were present in the gastrodermis of the body wall and strongly enriched in tentacles (Extended data fig. 1b, d) and in three different parts of the mesenteries: the intermediate tract (IT) (Fig. 1a, b-e), the somatic gonad epithelium (SGE) (Fig. 1a, b’-e’) and the trophic tract (TT) (Fig. 1a, b”-e”). In contrast, all other mesentery regions remain largely devoid of beads (Fig. 1a, b-e”). Phagocytosis of 1μm Fluospheres becomes apparent already after 30 minutes of incubation (Fig.1 b-b”) and leads to intracellular accumulation after 2 and 24 hours of incubation (Fig.1c-d”). A similar uptake distribution is observed using fluorescently labelled, heat-killed *E. coli*, confirming the physiological relevance of BSA-coated bead experiments (Extended data fig. 2a-a’’).

Interested to explore if endocytosed microspheres become mobilized in the tissue over time, we used a 4-hours incubation pulse of 20nm beads and found similar distributions after 20 hours, 7 days and 14 days-long chase periods (Extended data fig. 2b, c-e’’), indicating that no transport occurs between body regions. Bi-weekly incubations over four weeks led to an intracellular accumulation of 20nm nanobeads in the apical region of SGE cells (Extended data fig. 2f, g). Notably, a subset of mucus cells, abundant within the SGE^35^, shows no nanobead uptake (Extended data fig. 2f, g). Altogether, we found that three mesenterial regions, including the somatic gonad epithelium, have increased phago- and micropinocytosis activities.

Phagocytic cell types are well-studied on the molecular level in the context of immunity (e.g. macrophages) but not in the context of nutrition^36^. We therefore investigated the spatial expression profiles of endocytic pathway marker genes and transcription factors in nutritive phagocytic cells of the *Nematostella* mesenteries. Genes were selected for analysis based either on their value as conserved marker genes for cellular processes (e.g. phagocytosis, intracellular digestion), because of their presence in putative endocytic ‘metacells’ of a whole-polyp, single-cell sequencing dataset^37^, or as spatial reference to juvenile nutrient storage regions^38^. We found that *lipopolysaccharide binding protein/bactericidal permeability-increasing protein* (*lbp/bpi*, Extended data fig. 3a) and C-type lectin *mannose receptor* (*mannR*, Extended data fig. 3b) gene orthologs of *Nematostella* are specifically expressed in the endocytic TT, IT and SGE (Fig. 2a-a”, b-b”, j). Their bilaterian orthologs recognize bacterial lipopolysaccharide and glycans as well as soluble macromolecules and large particulate matter, and MannR is known to induce phagocytosis and clathrin-mediated endocytosis^39–41^. Their *Nematostella* expression thus supports a dual role for endocytic regions in both immunity and nutrition. Next, we checked the expression of *Nematostella* orthologs of *cdc42* and *rhoA* genes, playing key roles in phagocytic cup formation, and of *low-density lipoprotein-like* (*ldlr-like*), *clathrin light chain* (*clatLC*) and *heavy chain* (*clatHC*) genes, all encoding for proteins with key roles in clathrin-mediated endocytosis. Expression of all these genes overlaps with bead uptake in IT, TT and SGE regions (Fig. 2c-g’’, j), supporting phagocytosis and clathrin-mediated endocytosis as the main endocytic mechanisms there. Additionally, the predominant expression of gene orthologs encoding for glycosidases (*α-glucosidase, α-mannosidase*), proteases (*cathepsin*) and cholesterol transport/recycling (*npc2*) proteins, in all or a subset of the three endocytic mesenterial regions, supports an increased level of intracellular digestion (Extended data fig. 4). In juvenile *Nematostella*, the median, lipid-storing part of the mesentery was previously reported to co-express a combination of *foxC, six4/5* and *nkx3*/*bagpipe* transcription factor genes, revealing striking similarities to lateral mesoderm derivatives of bilaterians such as somatic gonad or nutrient storage tissues^38,42^. How this juvenile region relates to adult endocytic structures was yet to be determined. While nkx3/*bagpipe* expression levels were below detection limits, *foxC* and *six4/5* genes were expressed in all endocytic regions of the adult mesentery (Fig. 2h-j), indicating similarities in function and transcription factor profile to the lipid-uptaking region of juvenile mesenteries.

**Figure 2:**
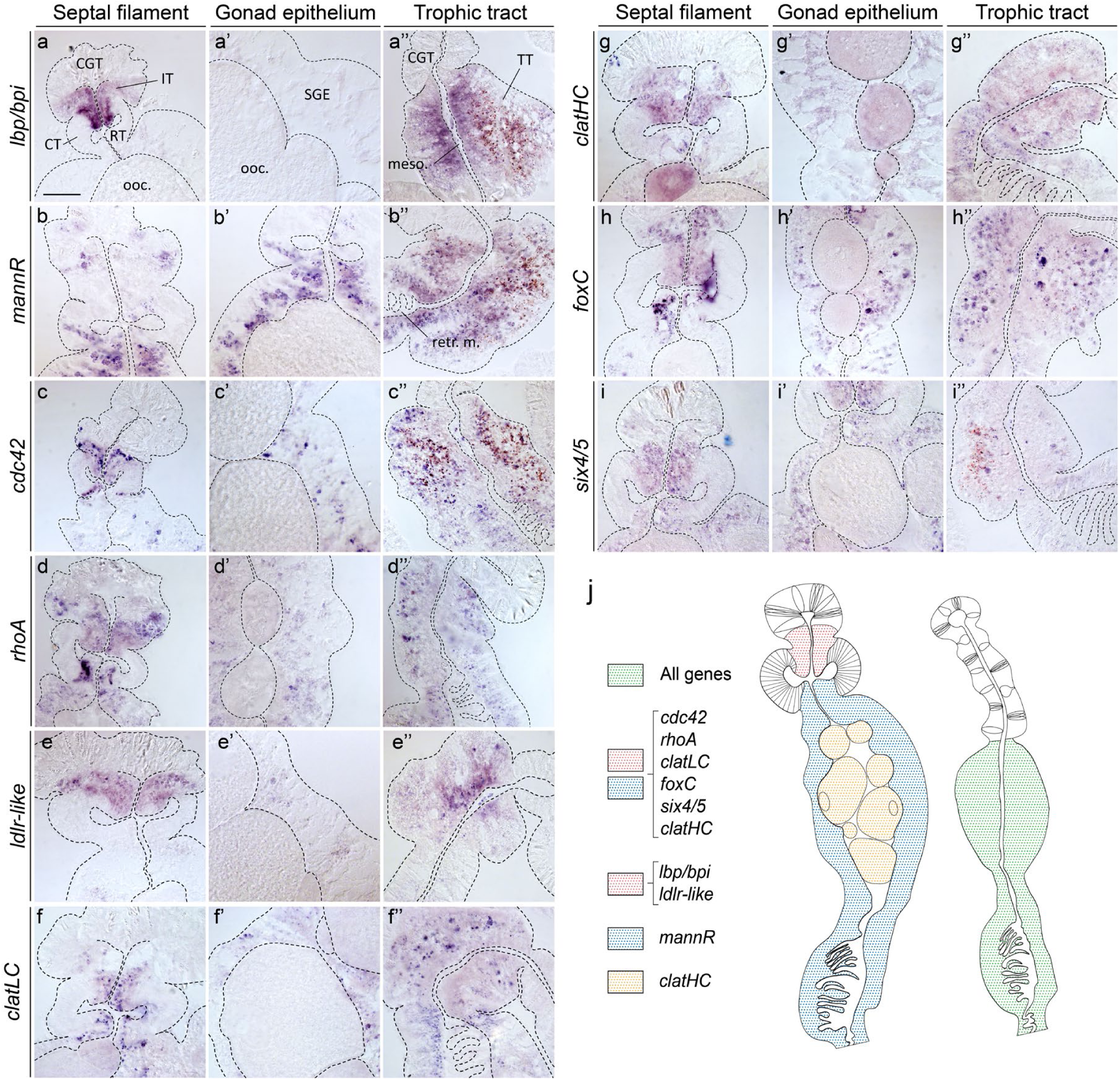
Phagocytosis and receptor-mediated endocytosis-related genes are expressed in endocytic regions of the mesenteries. (a-i”) mRNA expression patterns of *Nematostella* orthologs of the *lbp/bpi* (a-a’’), *mannR* (b-b’’), *cdc42* (c-c’’), *rhoA* (d-d’’), *ldlr-like* (e-e’’), *clatLC* (f-f’’), *clatHC* (g-g’’), *foxC* (h-h’’) and *six4/5* (i-i’’) genes in the septal filament (a-i), gonad epithelium (a-i’) and trophic tract (a’’-i’’). See text for gene descriptions. (j) Summary schematics of gene expression patterns. All mesenteries oriented with distal end to the top. All data figures represent cross sections of adult *N. vectensis* mesenteries stained by *in situ* hybridization. Scale bar: 50 μm. CT: ciliated tract, CGT: cnidoglandular tract, *clatHC* and *LC*: *clathrin heavy* and *light chain*, SGE: somatic gonad epithelium, IT: intermediate tract, *lbp/bpi*: *lipopolysaccharide binding protein/bactericidal permeability increasing protein, ldlr-like*: *low-density lipoprotein receptor-like gene, mannR*: *mannose receptor*, meso.: mesoglea, ooc.: oocyte, RT: reticulate tract, retr. m.: retractor muscle, TT: trophic tract.

Our characterization of cellular particle uptake modalities and gene expression profiles has highlighted the specialization of distinct mesenterial regions in food particle endocytosis. We next tested if these regions provide dietary nutrients to the oocytes by studying the cells and molecules mediating the transport of fatty acids from the gastric cavity into the oocytes.

### Dietary fatty acids reach the oocyte by trans-epithelial transport

Spatial detection of neutral lipid storage using Oil Red O staining confirms that lipids are a major yolk component (Extended data fig. 5a). It also shows high levels of neutral lipids largely overlapping with endocytic regions (SGE, IT and TT) in both males and females (Extended data fig. 5a-c). In contrast, lipid levels appear low in non-endocytic regions such as the CT or CGT, suggesting that only specialized endocytic cells and gametes possess lipid transport or storage abilities. We explored the dynamics of fatty acid uptake and transport from the gastric cavity towards the oocyte by developing a whole-body pulse-chase assay using a ‘clickable’, alkyne-modified oleic acid (alkyne-OA)^43^. Previous studies used alkyne-OA to reliably mimic oleic acid and study its metabolism in yeast, *Drosophila* and mammalian cells^43^. As oleic acid is abundantly found in anthozoan storage lipids, it is well-suited to track dietary fatty acids in *Nematostella*^44,45^. A 4-hours pulse/20-hours chase experiment using alkyne-OA-enriched brine shrimps resulted in labelling of the IT, the SGE (Fig.3 a, b, d) and the distal TT (Extended data fig. S6a, c), largely overlapping with endocytic regions. Notably, both IT and SGE become devoid of alkyne-OA after 7 days of chase (Fig. 3c, e), suggesting lipid catabolism or translocation. In contrast, alkyne-OA in TT cells shift location from the apical side of distal cells (at 20 hours chase; Extended data fig. 6a, c) to the basal side of proximal cells (at 7 days chase; Extended data fig. 6b, d) with small vesicles apparent in the mesoglea, possibly hinting at basolateral lipid secretion into the ECM (Extended data fig. 6d, arrowheads).

**Figure 3:**
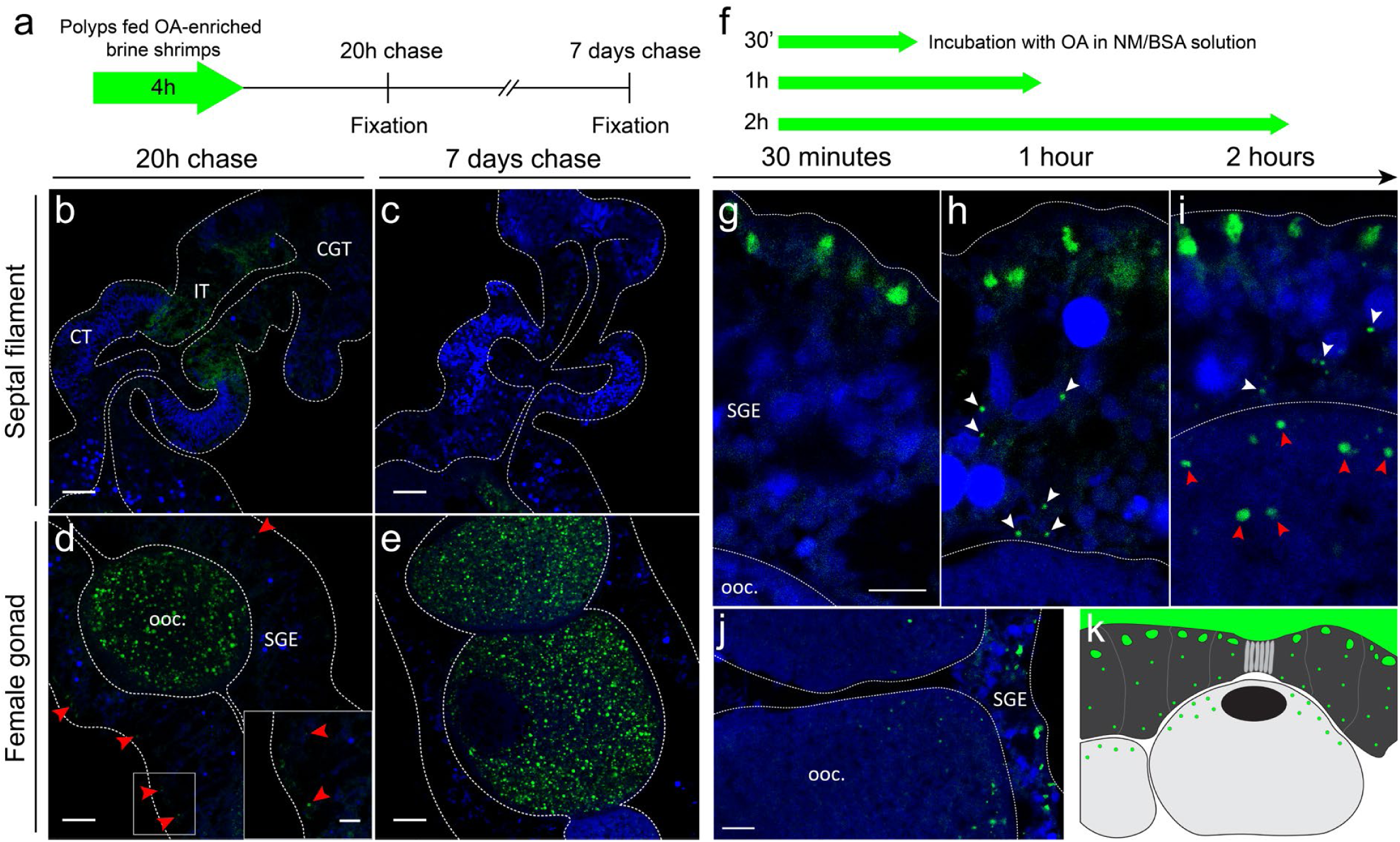
Trans-epithelial transport of alkyne-OA occurs from the gastric cavity into oocytes. (a-e) A pulse-chase experiment using a 4-hours pulse of alkyne-OA-enriched brine shrimps reveals long-term changes in lipid mobilization. In the IT, alkyne-oleic acid (green) is detected faintly after 20 hours (b) but not after 7 days (c) of chase. Similarly, small positive vesicles are present in the gonad epithelium after 20 hours (d and insert within, red arrowheads) but are absent after 7 days (e). Note that alkyne-OA is present in oocytes after both 20 hours and 7 days of chase (d, e). (f-k) Incubation with BSA-coupled alkyne-OA over short time periods highlights the timeline and dynamics of alkyne-OA uptake. Alkyne-OA is progressively detected at the apical surface of SGE cells after 30 minutes (g), their basal half after 1 hour (h, white arrowheads), and in oocytes after 2 hours of incubation (i, red arrowheads). Note that the signal in oocytes is restricted to the side of the cells directed towards the gonad epithelium (j). (k) Schematic summary of trans-epithelial alkyne-OA transport. Mesenteries oriented with distal end to the top (b-e, j) or left (g-i, k). All data figures show single confocal plane images of adult female *N. vectensis* mesentery cross sections. Blue: DAPI nuclear stain. Scale bars: (b-e) 25μm, insert in (d) and (g-j) 10 μm. CGT: cnidoglandular tract, CT: ciliated tract, IT: intermediate tract, OA: oleic acid, ooc.: oocyte, SGE: somatic gonad epithelium.

In oocytes, alkyne-OA strongly accumulates in vesicles after 20 hours and 7 days of chase (Fig. 3 d, e), confirming major dietary fatty acid movement from the gastric cavity into ECM-based oocytes. We increased the temporal control and resolution of alkyne-OA delivery by directly injecting alkyne-OA/BSA solution into the gastric cavity (Fig. 3f-j, Extended data fig. 7). This optimized delivery method yielded similar results to feeding enriched brine shrimps, which relied highly on individual rates of extracellular digestion (compare Extended data fig. 7b, c with Fig. 3d, e and Extended data fig. 7d, e with Extended data fig. 6a, b). We found that 30 minutes after delivery, alkyne-OA localizes to large apical vesicles in SGE cells (Fig. 3g). Within 1 hour of incubation, smaller vesicles appear in median and basal regions of SGE cells suggesting fast, intracellular, apical-to-basal transport (Fig. 3g, h). Between 1 and 2 hours of delivery, alkyne-OA vesicles start accumulating intracellularly at the periphery of the oocyte adjacent to the SGE (Fig. 3i-k). These observations are in full agreement with a trans-epithelial transport of alkyne-OA from the gastric cavity through the SGE into oocytes via the surrounding ECM. Our inability to detect alkyne-OA in the ECM speaks for a rapid clearance by high levels of endocytosis into oocytes, strongly supported by ultrastructural observations^27^. Notably, our experiments did not reveal any alkyne-OA or fluorescent bead accumulation in trophonema cells of the female SGE, despite this structure being previously proposed as the main nutrient shuttling cells during vitellogenesis in *Nematostella* and other sea anemones^24,26,46^. As our data challenges the historical hypothesis for nutrient transport into the oocytes during vitellogenesis, we next investigated how SGE-ECM lipid transport is mediated on the molecular level.

To do so, we studied the expression of the single *Nematostella* orthologs of *vtg* and *apoB* apolipoprotein genes (Extended data fig. 8) and three *Nematostella vldl receptor* paralogs conserved in other sea anemones and corals (Extended data fig. 9). We confirm previous reports from *Nematostella* and other anthozoans that the SGE expresses high levels of *vtg* transcripts (Fig.4 a-a”)^47–50^. In contrast, no *vtg* transcript was detected in oocytes despite Vtg protein representing approx. 60% of the protein content of the mature egg in *Nematostella*^51^. This result indicates that endocytosis contributes to a large extent to Vtg protein accumulation during vitellogenesis in anthozoans. *vldlr-A1, -A2* and *-B* genes (Extended data fig. 9) are all expressed in growing oocytes of different sizes, supporting a role in receptor-mediated apolipoprotein uptake during vitellogenesis (Fig. 4b-e). *vldlr-A1* and *vldlr-A2* are additionally expressed in uncharacterized cells of the CT and reticulate tracts of the septal filament (Fig. 4b”, c”). As Vtg constitutes the most abundant protein in mature oocytes, our data strongly suggest that Vtg mediates lipid transport from SGE cells into growing oocytes by VLDLR-mediated endocytosis, as widely found among bilaterians^10,52^.

**Figure 4:**
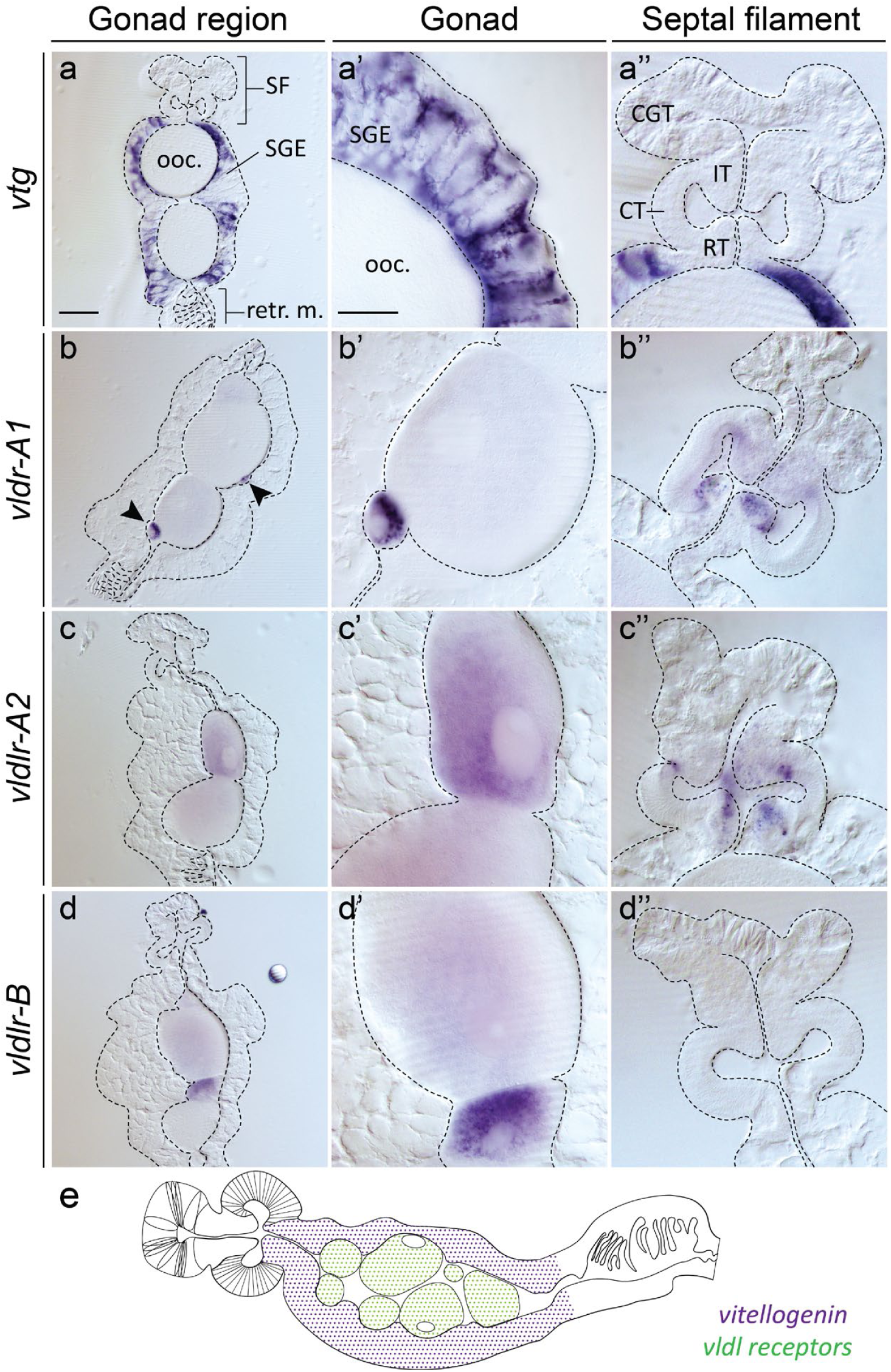
Complementary expression of *Nematostella vtg* and *vldlr* gene orthologs supports lipid transport from the SGE into oocytes. (a-d”) mRNA expression pattern of *vtg* in somatic gonad epithelium (a-a”) and its potentially conserved lipoprotein receptors *vldlr-A1* (b-b”, small eggs marked by arrowheads), *vldlr*-*A2* (c-c”), and *vldlr*-*B* (d-d”) in oocytes of different sizes (b-d, b’-d’) and partly in the septal filament (b”-d”). (e) Schematic summary of the complementarity of *vtg*/*vldlr* expression patterns. Mesenteries oriented with distal end towards top (a-d’’) or left (e). All data figures show cross sections of adult female *N. vectensis* mesenteries (gonad region). Scale bars: (a-d) 100μm, (a’-d’’) 50μm. CT: ciliated tract, CGT: cnidoglandular tract, SGE: somatic gonad epithelium, IT: intermediate tract, ooc.: oocyte, RT: reticulate tract, *vldlr*: *very low-density lipoprotein receptor, vtg*: *vitellogenin*.

Although ApoB proteins are found in bilaterians, cnidarians, placozoans (Extended data fig. 8), their expression profile and role in lipid transport have not been characterized in any non-bilaterian animal. In *Nematostella, apoB* expression colocalizes with *vtg* expression in the SGE but extends also to the IT and TT, thus overlapping with endocytic regions (Fig. 5a-c). This result suggests a more systemic role for ApoB in lipid transport, as usually found in bilaterians^1^. To further explore this possibility, we aimed to study if ApoB proteins and alkyne-OA label colocalize. We therefore generated a transgenic reporter line using CRISPR/Cas9-mediated knock-in technology resulting in a genomically-encoded, C-terminal ApoB-PSmOrange fusion protein. As expected, immunofluorescence detection shows the colocalization of ApoB-PSmOrange protein to endocytic regions with high *apoB* transcript levels (compare Fig. 5a-c and d-i). ApoB-PSmOrange localizes to larger vesicles throughout the cells of the IT (Fig. 5d, e) and male SGE (Fig. 5h), while these vesicles accumulate in the apical regions of the TT and female SGE cells (Fig. 5g, f, i). Growing oocytes were devoid of signal (Fig. 5g), as supported by low levels of ApoB protein detected by mass spectrometry in spawned eggs^51^. Strikingly, however, ApoB-PSmOrange protein was detected in the spermaries, with higher intensities in peripheral, immature cells compared to central, mature sperm cells (Fig. 5h). While we found broad overlap between cells with high ApoB-PSmOrange protein and alkyne-OA uptake levels, we found only few instances of intracellular colocalization in the SGE after a 2-hours alkyne-OA pulse (Fig. 6a-b’’). Interested to investigate a potential role for ApoB in systemic lipid transport, we looked at the distribution of ApoB-PSmOrange and alkyne-OA after a 4-hours alkyne-OA incubation pulse followed by a 7-days chase. Strikingly, we observed that the two signals colocalize in mesogleal cells of the trophic region (Fig. 6 c-d’’). This is consistent with previous observations of nutrient transport by motile mesogleal cells, the amoebocytes^53–55^. In the spermaries, ApoB-PSmOrange and alkyne-OA localize to adjacent, but distinct vesicles (Fig. 6e-e’’). Overall, our data supports a potential function of ApoB in systemic lipid transport via the mesoglea and during spermatogenesis, but not during oogenesis.

**Figure 5:**
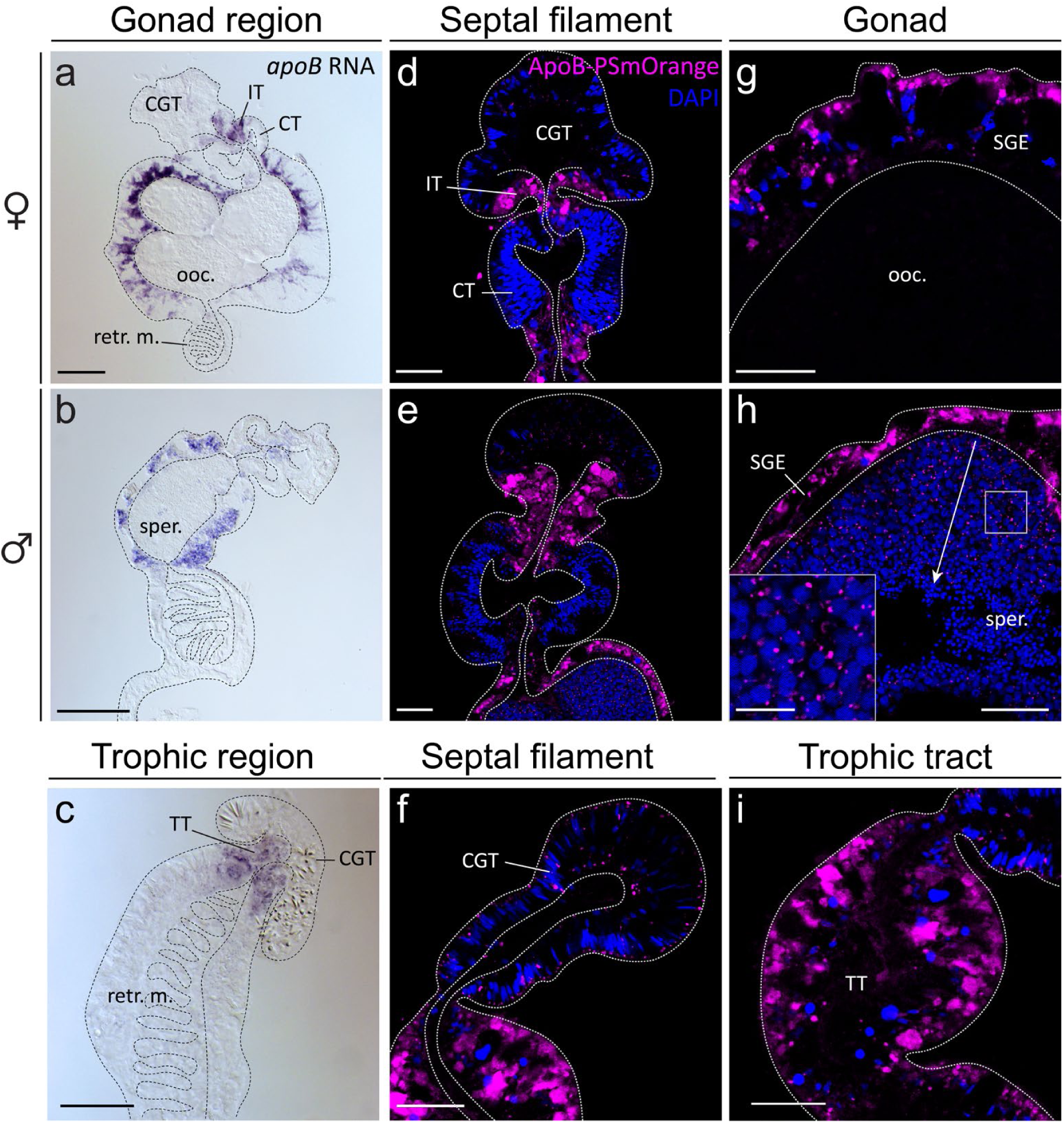
*Nematostella apoB* mRNA and ApoB-PSmOrange fusion protein localisation suggests a role for ApoB in spermatogenesis but not in vitellogenesis. (a-c) *apoB* mRNA expression in the gonad region of female (a) and male (b) polyps, and in the trophic region (c). (d-i) ApoB-PSmOrange protein localization (magenta) is consistent with *apoB in situ* hybridization data (a-c) in the TT and in the IT and SGE of the female and male gonad region. Notably, ApoB-PSmOrange is found in spermaries (h and close-up within). A peripheral to central gradient of decreasing signal intensity coincides with the progression of sperm cells maturation stages (arrow). Mesenteries oriented with distal end to the top (a-f, i) or left (g, h). Cross sections of adult *Nematostella* mesenteries. Blue: DAPI nuclear stain. All fluorescent images represent single confocal planes. Scale bars: (a-c) 100μm, (d-i) 25μm, close up in (h): 10μm. CT: ciliated tract, CGT: cnidoglandular tract, *clatHC* and *LC*: *clathrin heavy* and *light chain*, SGE: somatic gonad epithelium, IT: intermediate tract, ooc.: oocyte, retr. m.: retractor muscle, sper.: spermary, TT: trophic tract.

**Figure 6:**
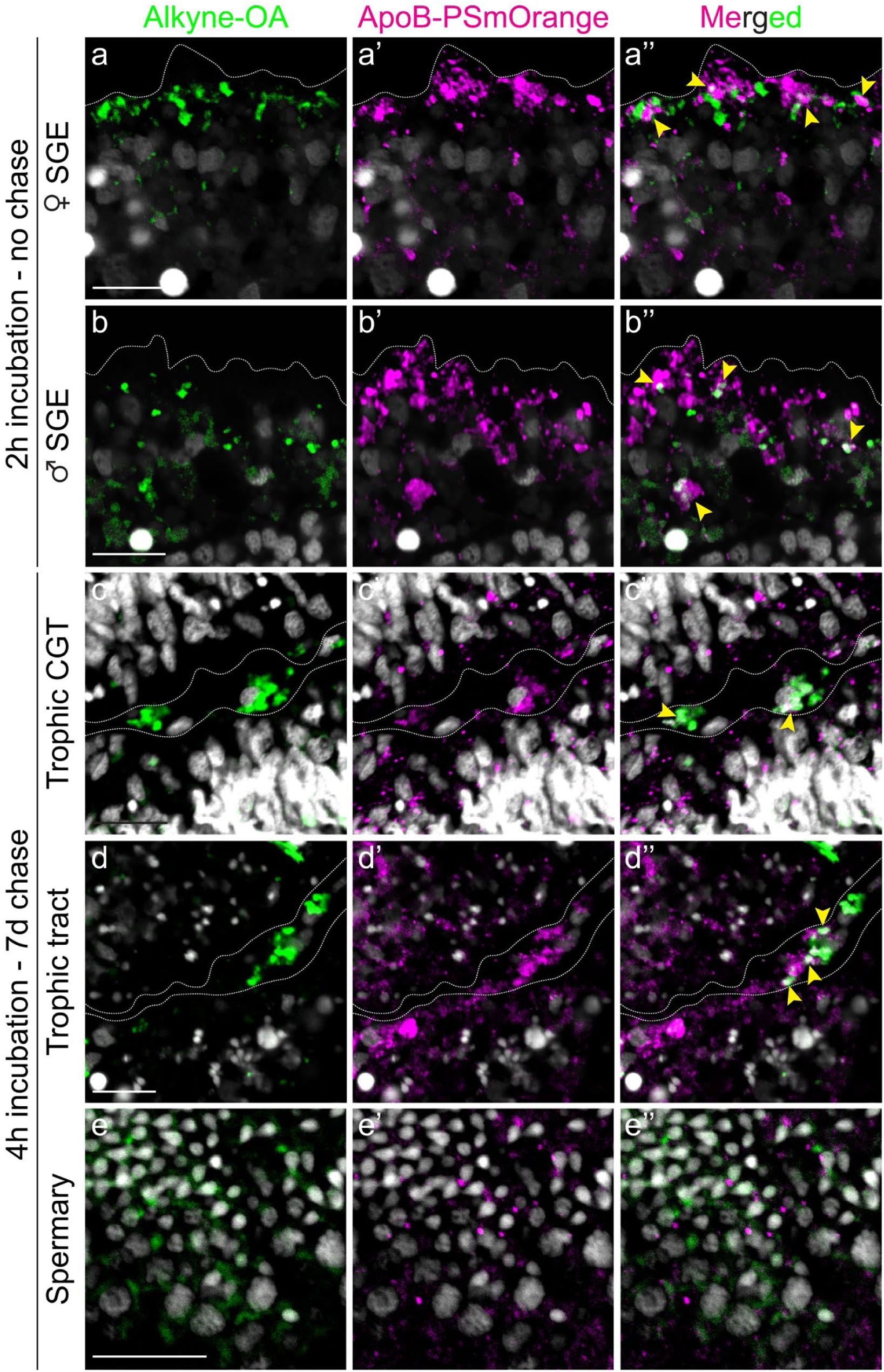
Colocalization of alkyne-oleic acid and ApoB-PSmOrange in a subset of SGE vesicles and in mesogleal cells of the trophic region. Co-staining of alkyne-OA (green) and ApoB-PSmOrange (magenta) after 2 hours alkyne-OA pulse (a-b’’) or 4 hours pulse and 7 days chase (c-e’’). Only few intracellular vesicles (a’’,b’’ arrowheads) show colocalization of alkyne-OA with ApoB-PSmOrange in the female (a-a’’) or male SGE (b-b’’). Interestingly, mesogleal cells with alkyne-OA and ApoB-PSmOrange colocalization are found in the trophic cnidoglandular tract (c-c’’, arrowheads) and trophic tract (d-d’’, arrowheads). Alkyne-OA and ApoB-PSmOrange vesicles locate adjacently but do not appear to overlap in the spermaries (e-e’’). Mesenteries oriented with distal end to the left. All images represent single confocal plane images of adult *Nematostella* mesenteries cross sections. White: DAPI nuclear stain. Scale bars: 10μm. CGT: cnidoglandular tract, SGE: somatic gonad epithelium.

## Discussion

In most bilaterian animals, dietary nutrients travel from the gut epithelium to oocytes via ECM-based circulatory systems or, more rarely, through coelomic cavity fluids^10,56,57^. Due to the lack of data from non-bilaterian animals, the emergence of systemic nutrient transport and their underlying cellular and molecular features during animal evolution remains highly speculative. We have therefore retraced the cellular path of dietary nutrients, more specifically lipids, from the gastric cavity to the oocytes during vitellogenesis in the sea anemone *Nematostella vectensis*. We have shown that the gastrodermal epithelium, although it is in direct contact with food particles and nutrients, does not show a uniform distribution of phagocytosis, micropinocytosis and lysosomal activities. Instead, our analyses of bead uptake assays and marker gene expression show that these activities are enriched in the tentacle gastrodermis as well as the IT, SGE and TT mesenterial regions. In corals and symbiotic sea anemones (e.g. *Exaiptasia*), carbon-fixing *Symbiodinium sp*. dinoflagellates locate mainly to the tentacle gastrodermis and the IT and SGE regions^25,58^. These parallels confirm the importance of these regions in nutrient acquisition in sea anemones and corals and raise questions about symbionts’ ability to permanently populate these highly endocytic regions in some anthozoan species but not in others.

As our molecular knowledge of animal phagocytosis relies almost entirely on the study of innate immunity cell types (e.g. macrophages), we provide one of the first molecular characterization of animal trophic phagocytes^36^. Notably, the set of genes found expressed in nutritive endocytic regions in *Nematostella*, composed of immunity-related pattern recognition proteins (LPS/BPI, C-type lectins), phagocytic cup and clathrin-mediated endocytosis components as well as lysosomal enzymes and transporters, is largely shared with immune cells in vertebrates and flies^59–62^. This raises the possibility that *Nematostella* trophic phagocytes are bi-functional and carry out nutritive as well as immunity-related functions, e.g. in controlling the microbiome composition or symbiont uptake in the gastric cavity. Currently, however, a reconstruction of the evolution of trophic and immune phagocytic cell types remains difficult due to the lack of comparative molecular data from bilaterian trophic phagocytes^36^. Nevertheless, the expression of *foxC and six4/*5 transcription factors in *Nematostella* endocytic regions is reminiscent of the lateral mesoderm regions of bilaterians, which typical derivatives include blood immune cells, nutrient storage tissues (e.g. insect fat body) or somatic gonad tissue^63^.

We identified the SGE among mesenterial endocytic regions as highly relevant for dietary nutrient transport during vitellogenesis. Indeed, our data proposes that the SGE play a key role in dietary particle and lipid uptake as well as a Vitellogenin-producing tissue. Our alkyne-oleic acid tracking experiments strongly suggest that dietary fatty acids are rapidly absorbed and transported through the SGE into the ECM before getting internalized via endocytosis by oocytes, as corroborated by ultrastructural observations^27^. As Vtg is highly expressed in the SGE and highly abundant in the mature egg, we propose that it constitutes the key lipid transporter from the SGE through the ECM into oocytes during vitellogenesis in *Nematostella*. We found three *Nematostella vldl receptor* paralogs expressed during oocyte growth, of which VLDLR-B (Uniprot: A7RXU8) was previously identified in the proteome of mature eggs^51^. This result supports that Vtg endocytosis is mediated by VLDL receptors in this species. Together, our data suggests that a Vtg ligand/VLDL receptor pair has been evolutionary conserved during vitellogenic lipid transport since the last common bilaterian-cnidarian ancestor. ApoB protein, involved in systemic lipid transport in bilaterians, appears to have no direct role during vitellogenesis in *Nematostella*. Although abundantly found in the SGE, it was detected at low levels in oocytes using mass spectrometry^51^, and below detection levels using immunofluorescence. Instead, high ApoB-PSmOrange protein levels detected in all endocytic tissues and partially colocalizing with apical alkyne-OA suggest potential roles for ApoB in fatty acid uptake and intracellular transport. Strikingly, we found ApoB protein strongly colocalizing with alkyne-OA in single cells in the mesoglea (ECM), suggesting systemic lipid transport by mesogleal cells, such as the amoebocytes^53–55^. In addition, we find a potential role for ApoB in lipid transport or metabolism during spermatogenesis that requires further investigation.

A prevalent hypothesis about the origin of circulatory systems in bilaterians proposes that they evolved from a simple ECM-based nutrient transport system^20^, but this model lacked experimental support until now. Here, we show the first evidence that Vtg/ApoB- and VLDLR-mediated lipid transport through the extracellular matrix space is not only conserved in bilaterians, but also found in a sea anemone. This suggests that nutrient transport through the extracellular space has evolved early during animal evolution, and that the spatial separation between food uptake and metabolism served as a mechanistic basis for the evolution of intricate ECM-based circulatory systems such as the hemolymph or blood vascular systems in insects and vertebrates.

## Acknowledgements

We thank E. Hambleton for kindly providing plasmids encoding *npc2* genes, and all Steinmetz lab members for helpful discussions and feedback on the manuscript.

## Materials and Methods

### Animal culture

The *Nematostella vectensis* polyps are derived from the original culture established by C. Hand and K. Uhlinger^64^. Adult animals are maintained in the dark at 18°C in 16ppm diluted sea water (Nematostella medium, NM) and spawning is induced approximately every three weeks by a shift in light and temperature conditions as described previously^65^.

### Particle uptake assays and fixation

Fluorescent latex beads of 1μm (Thermo Fischer, carboxylate-modified, crimson fluorescent Ex/Em = 625/645) and 20nm diameter (Thermo Fisher, carboxylate-modified, yellow-green fluorescent Ex/Em = 505/515) were coated in BSA for 2 hours in 2% BSA/NM solution at a 1:1 ratio and further diluted 1:100 in 0.1M MgCl2/NM before being fed to animals.

For the bacteria uptake assay E. coli K-12 strain Bioparticles (Thermo Fisher, Alexa Fluor 488 conjugate Ex/Em = 495/519) were resuspended in filtered water to reach a stock concentration of 3×108 particles/ml and further diluted to 1:100 in 0.1M MgCl2/NM before use. No BSA coating was applied.

For all particle uptake assays, animals were initially incubated in 0.1M MgCl2/NM for 20 minutes to avoid muscle contraction and kept in this medium for the duration of the incubation time when it did not exceed 4 hours. For longer incubations, animals were kept in MgCl2/NM for the first 4 hours then transferred to fresh NM. The fluospheres or E. coli solution was injected inside the body cavity through the mouth using a thinned glass Pasteur pipet and the animals were kept in the dark at 18°C for the duration of the experiment.

At the end of the assays, animals kept in NM were transferred back to 0.1M MgCl2/NM solution and left to relax for 20 minutes. The body cavity of the animals was flushed with this medium through the mouth to ensure full extension of the mesenteries. The polyps were then transferred to a 3.7% Formaldehyde/NM solution and opened longitudinally along the body column to ensure penetrance of the fixative and to allow non-internalized particles to be washed off. The fixation was conducted overnight at 4°C, after which the mesenteries were dissected in fixative and cut in 3-5mm long pieces. The tissue pieces were washed in 1x PBS/0.1% Tween-20, counterstained with Alexa-488 or Alexa-647 phalloidin (Thermo Fisher) and DAPI nuclear staining before being processed for cryosectioning.

### Identification of candidate genes and orthology

Candidate genes were identified using in-depth literature searches, published *N. vectensis* egg proteome^51^ and transcriptome^37^ datasets as well as BLAST of the publicly available genome^66,67^. Orthology was confirmed by reciprocal pBLAST using the NCBI BLAST platform (http://www.ncbi.nlm.nih.gov/blast/) and phylogenetic analyses.

### Phylogeny

Amino acid sequences were retrieved using the NCBI BLAST platform and aligned using MUSCLE v3.8.31^68^. Sequence stripping with GBlocks^69^ using the least conservative parameters (Min. Num. of Seq. for Flank Pos.: lowest possible; Min. Block Length: 2; Gaps set to “half”). Stripped alignments were tested for the best fitting maximum likelihood parameters (LDLR, LRP1: WAG+I+G4; Vtg: LG+I+G4) and maximum likelihood trees calculated using iqTree using ultrafast bootstrapping (1000 replicates). Bayesian posterior probabilities calculated using MrBayes 3.2.7^70–72^ using LG+I+G8 (Vtg) or WAG+I+G4 (LDLR, LRP1) model parameters, two parallel runs, a temperature of 0.2 (LDLR, LRP1) or 0.05 (Vtg), eight chains, 1 (LDLR, LPR1) or 2 (Vtg) swaps tried at each swapping generation, and a sample frequency of every 10 generations. Convergence was reached after 20.000 (LDLR), 80.000 (Vtg) or 100.000 (LPR1) generations. A consensus tree was calculated using a burn-in of 5000 generations. Maximum-likelihood phylogenetic analyses were run on the IQ-TREE web server (http://iqtree.cibiv.univie.ac.at/, Trifinopoulos et al. 2016) using the best-fit substitution model and an ultrafast bootstrap analysis with 1000 alignments. Trees were visualized using FigTree v1.4.4 (https://github.com/rambaut/figtree/releases) and modified using Adobe Illustrator 2022 for MacOS.

### Gene cloning and RNA probe synthesis

RNA was extracted from whole adult female Nematostella using Trizol (Thermo Fisher), following the manufacturer’s protocol. cDNA was obtained using the SuperScript III first-strand synthesis system (Thermo Fisher). Fragments of the genes of interest were amplified using primers designed with the Primer3Plus online tool (http://www.bioinformatics.nl/cgi-bin/primer3plus/primer3plus.cgi). The fragments were then inserted in a pGEM-T Easy vector (Promega) and transformed into One Shot Top 10 chemically competent *E. coli* (Invitrogen). Cloned sequences were verified by Sanger sequencing at the sequencing facility of the Department of Biological Sciences, University of Bergen, Bergen, Norway. Antisense RNA probes were generated using a T7 or SP6 MEGAscript Kit (Invitrogen) and labeled with DIG RNA Labeling Mix (Roche) using previously published protocols^74^.

### *In situ* hybridization

Adult polyps were fixed following the protocol described above. The dissected pieces of mesenteries were washed in 100% methanol until all pigmentation was removed and subsequently stored in 100% methanol at −20°C.

The urea-based in situ hybridization protocol was adapted from Sinigaglia et al. 2018, with the following parameters and changes. After progressive rehydration in MeOH/PTx (0.3% Triton X-100 in 1xPBS pH 7.4), the tissue was digested with Proteinase K 2.5μg/ml for 5 minutes at room temperature. An additional fixation step (2 min in 0.2% glutaraldehyde/PTx followed by 1 hour in 3.7% formaldehyde/PTx) was added before overnight blocking in a hybridization mix consisting of 50% 8M urea, 5x SSC pH 4.5, 0.3% Triton X-100, 1% SDS, 100 μg/ml heparin and 5 mg/ml Torula yeast RNA. Background staining could be reduced by adding 5% dextran sulfate (MW > 500,000, Sigma-Aldrich) and 3% Blocking Reagent (Roche) to the hybridization mix during overnight blocking and hybridization of the probe. Hybridization was conducted over two days. The probe concentration was 0.75 ng/μl. Stringent washes varied between 2x SSC/0.3% Triton X-100 (SSCT) and 0.1x SSCT. After the SCCT washes, the tissue pieces were washed with 1x PBS/0.1%BSA/0.3% Triton X-100.

The staining reaction (NBT 4.5μl/ml and BCIP 3.5μl/ml in AP buffer) was performed for approximately 5 minutes for strongly expressed genes to up to 8 hours for weakly expressed genes.

### Oleic acid assays

Stock solution of alkyne-modified oleic acid (17-yne) (Avanti Polar Lipids, ethanol solution 1mg/ml) was aliquoted, and the ethanol evaporated in a vacuum concentrator.

For pulse-chase experiments in Fig. 3a-e and Extended data Fig. 6, the fatty acids were resuspended in DMSO to a concentration of 3mg/ml, and further diluted to the final concentration of 0.3mg/ml in sea water containing freshly hatched *Artemia salina* nauplii. The artemia were left to incubate in this solution for 2 hours, after what they were washed several times with NM on a fine mesh filter and resuspended in NM. Overnight-fasted adult *Nematostella* polyps were fed a few drops each of the enriched artemia and left to catch and ingest the prey for 4 hours. They were placed in clean medium after the feeding period and fixed after a 20 hours or 7 days chase. During the 7 days chase, animals were fed daily with regular artemia.

For the oocyte uptake timeline experiment in Fig.3g-j, and for the experiments in Fig. 6 and Extended data Fig. 7, the fatty acids were directly resuspended in a 4 mg/ml fatty acid-free BSA/NM solution following the manufacturer’s protocol. Overnight-fasted adult *Nematostella* polyps were incubated in 0.1M MgCl2/NM for 20 minutes to avoid muscle contraction, after what the OA solution was injected in the body cavity through the mouth using a thinned Pasteur pipet. The animals were fixed 30 minutes, 1 hour or 2 hours after injection.

At the time of fixation, animals kept in NM were transferred to 0.1M MgCl2/NM solution and left to relax for 20 minutes while animals that had been incubated in MgCl2 during the assay were processed directly. Fixation was conducted following the same protocol as described above.

### Click-it reaction

In order to visualize the alkyne-modified oleic acid in the tissues of the animals, we performed a click-it reaction using an azide-modified fluorophore (Alexa Fluor 488 Azide, Invitrogen, Ex/Em = 495/519).

Fixed tissue pieces were blocked for 30 minutes at 4°C in a 3% BSA/1x PBS solution, followed by a 30 minutes permeabilization step at 4°C in PTx (0.3% Triton X-100 in 1xPBS pH 7.4). The click-it reaction was performed using the reagents of a Click-iT™ EdU Cell Proliferation Kit for Imaging (Invitrogen) following the manufacturer’s protocol. The staining reaction was conducted for 30 minutes at room temperature. The tissue was then washed in 1x PBS three times over a period of 2 hours to remove residual fluorophore, stained with DAPI nuclear staining before being processed for cryosectioning.

### CRISPR-Cas9 mediated generation of apoB-PSmOrange knock-in line

Two guide RNA (gRNA) target regions with putative cutting sites located 52bp upstream or 24bp downstream of the Stop codon of the Nematostella ApoB (v1g84136) were designed using CRISPOR^76^. Templates for gRNAs were generated using annealed and PCR-amplified oligos^77^: two T7- and gRNA-encoding oligo (Thermo Fisher, desalted): 5’-GAAATTAATACGACTCACTATAGTCCTGTGTACATGGATACGTGTTTTAGAGCTAGAAATAGCAAG-3’ (upstream of Stop codon); 5’-GAAATTAATACGACTCACTATAGactaatcctaattaccaagtGTTTTAGAGCTAGAAATAGCA AG-3’ (downstream of Stop codon); invariant reverse primer (Thermo Fisher, desalted): 5’-AAAAGCACCGACTCGGTGCCACTTTTTCAAGTTG ATAACGGACTAGCCTTATTTTAACTTGCTATTTCTAGCTCTAAAAC-3’.

Guide RNAs were in vitro transcribed using a T7 MegaScript transcription kit (ThermoFisher) followed by ammonium chloride precipitation and diluted in nuclease-free H2O to a final concentration of 1,5μg/μl. The DNA donor fragment for homology-mediated repair (HDR) consisted of a GGGGS2 linker and a PSmOrange ORF cloned in frame to the 3’ end of the apoB ORF, and framed by 1030bp long ‘left’ homology arm and a 973bp ‘right’ homology arm. pH2B-PSmOrange was a gift from Vladislav Verkhusha (Addgene plasmid # 31920; Subach et al. 2011). The entire construct was cloned into a pJet1.2 plasmid backbone (Thermo Fisher) using Gibson assembly master mix (NEB)^79^. Donor DNA fragment was PCR-amplified using 5’-biotin-labelled oligos flanking the donor fragment to increase integration efficiency as previously described^80^. Oligo sequences (Thermo Fisher; desalted) are: Forward oligo 5’-[Biotin]-TACGACT CACTATAGGGAGAGCGGC-3’; reverse oligo 5’-CCATGGCAGCTGAGAATA TTGTAGGA-3’.

The Cas9-mediated knock-in injection was performed by modifying previously described protocols for CRISPR/Cas9-mediate mutagenesis and used 0,75μg/μl nls-Cas9 protein (PacBio), 75ng/μl of each guide RNA, 70ng/μl column-purified donor DNA, and modified injection buffer containing 220mM KCl^81,82^. Out of approx. 2300 injected zygotes, one single ubiquitously fluorescent polyp has been identified and raised as founder animal. The success of the knock-in has been validated on the transcript level by in vitro synthesis of cDNA based on extracted total RNA from a mix of fluorescent F1 juvenile polyps. Based on this cDNA, the transitions between the apoB ORF and PSmOrange, and between PSmOrange and the 3’UTR have been amplified using oligos outside of the ‘homology arm’ regions. Sanger sequencing of gel-eluted and column-purified PCR fragments confirmed the flawless integration of the GGGGS2-PSmOrange fragment directly downstream of the ApoB open reading frame. Oligos used for PCR & sequencing: ApoB-Forward : 5’-GACTGCTAAGGACATGAAGCACTGC-3’; ApoB-Reverse (not used for sequencing): 5’ GACTTCGTCATACTCCGATACTGGAC-3’; PSmOrange-Forward (used with ApoB-Reverse): 5’-CAACGAGGACTACACCATCGTGG-3’; PSmOrange-Reverse (used with ApoB-Forward): 5’-CTCGAACTCGTGGCCGTTCAC-3’. Additional sequencing oligo: ApoB-Reverse-2: AATCATAGCACCACCTCACTGCG.

### Immunofluorescence

For the ApoB-PSmOrange knock-in reporter line immunofluorescence, animals were fixed overnight and dissected in 1xPBS containing 3.7% Formaldehyde, 0,5% DMSO and 0,1% Tween20 and stored in 100% methanol at −20°C before use. After progressive rehydration in 1xPBS, tissue pieces were blocked in 1xPBS containing 5% NGS/1% BSA/0.2% Triton X-100 for 30 minutes at room temperature. They were then incubated with primary antibody (rabbit anti-DsRed, 1:100, Takara Bio Clontech, order nr. 632496) overnight at 4°C. On the next day, the tissue was washed 6x with PTx, blocked again (see above), and incubated in the secondary antibody (goat-anti-rabbit-Alexa568, LifeTech, order nr. A11011). Finally, tissue pieces were washed several times and overnight in PTx, stained with DAPI nuclear staining and processed for cryosectioning.

### Cryotome sectioning

After overnight infiltration in 1x PBS/25% sucrose/20% OCT (Tissue-Tek O.C.T. Compound, Sakura) samples were mounted in 80% OCT/PBS medium, oriented for transversal sections and frozen on a metal block cooled with liquid N2. Sectioning was performed at −25 °C in a Leica Cryostat CM1850 and the 12μm sections were mounted on Thermo Scientific SuperFrost Plus adhesion slides (Thermo Fisher). After drying, the sections were post-fixed for 10 min in 3.7% FA/PBS, rinsed in 1xPBS, and mounted in glycerol.

### Oil Red O staining

Animals were fixed and dissected as described above for the alkyne-OA assay, and mesentery pieces were sectioned with a cryotome and post-fixed as described above. Oil Red O (ORO) stock solution was prepared by diluting 5mg/ml ORO powder in isopropanol, shaking at room temperature for 4 hours. The stock solution was further diluted to 60% in milliQ water, left shaking at room temperature for another 2 hours and filtered using a 0.22μm syringe filter to produce a working solution ^83,84^. The mesentery sections were incubated for 15 minutes with the ORO working solution, then washed in 1xPBS until the solution remained clear. Finally, the slides were mounted in glycerol before imaging.

### Transmitted light and confocal imaging

Images of in situ hybridization-stained tissue were taken on a Nikon Eclipse E800 using a 20x lens or 40x and 60x oil-immersion objectives. Images of fluorescent assays were taken either on a Leica SP5 confocal microscope using standard PMT detectors and a 20x or 63x oil-immersion lens, on a Leica SP8 confocal microscope using HyD detectors and 40x water-immersion or 100x oil-immersion objectives, or on an Olympus FLUOVIEW FV3000 confocal microscope using standard PMT detectors and a 60x silicon oil immersion lens. Transmitted light images were corrected for levels and color balance and cropped using Photoshop CC. Fluorescent stacks were processed, cropped and the levels corrected using Fiji.

## Notes

### Competing Interest Statement

The authors have declared no competing interest.

