## Supplementary data for "Evolutionary conserved aspects of animal nutrient uptake and transport in sea anemone vitellogenesis"

### Extended data figures

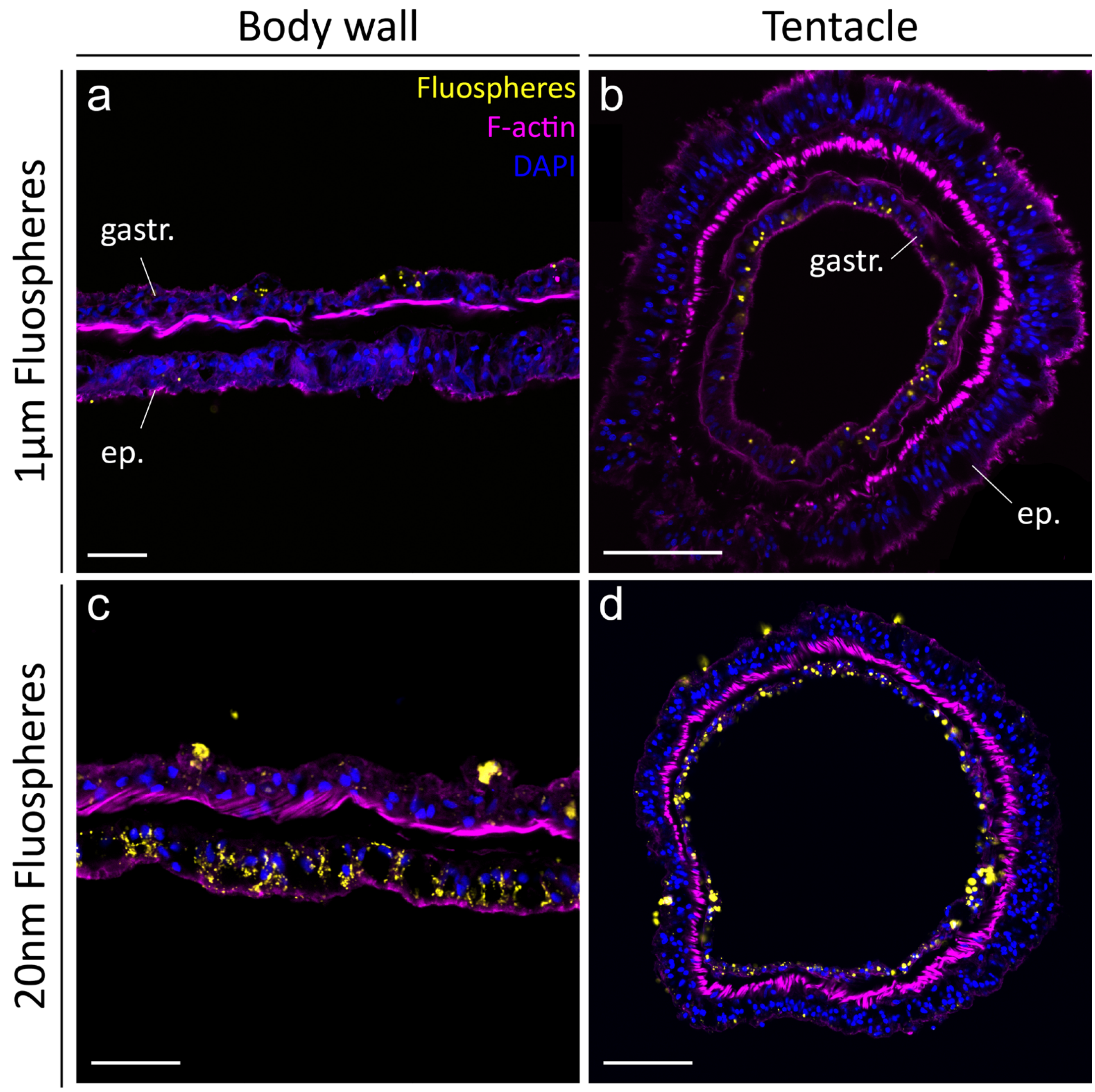


**Extended data figure 1: Differential uptake of 1µm and 20nm Fluospheres by epi- and gastrodermal epithelia of the body wall and tentacles.** Single plane confocal images of adult *Nematostella* body wall and tentacle after a 24-hours incubation with 1µm (a, b) or 20nm (c, d) Fluospheres. 1µm beads are taken up in relatively small numbers in the gastrodermis (a) while 20nm beads enriched in body wall epidermis (c). Both Fluospheres sizes are taken up by tentacle gastrodermis (b, d). Scale bars: (a, c) 25µm, (b, d) 50µm. ep.: epidermis, gastr.: gastrodermis.

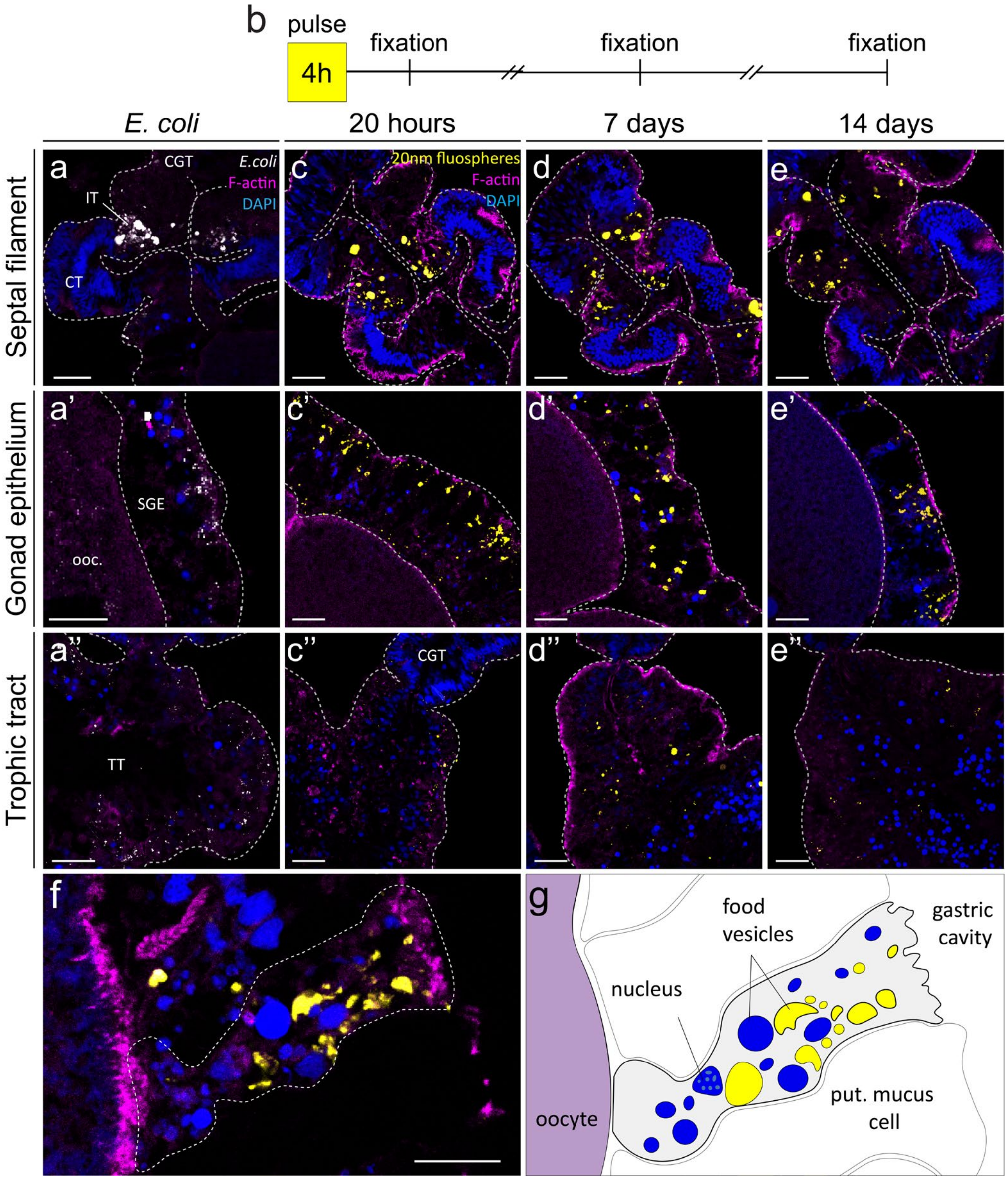


**Extended data figure 2: Distribution of fluorescent, inactivated *E. coli* bacteria and long-chased 20nm Fluospheres in *Nematostella* mesenteries.** (a-a'') A 4-hours incubation with fluorescent, inactivated *E. coli* bacteria reveals a distribution of uptake similar to 1μm and 20nm Fluospheres (see Fig. 1). (b-e'') A pulse-chase assay (experimental setup in (b)) shows the cellular retention and absence of transport of 20nm Fluospheres (yellow) between tissues after a 4 hours incubation followed by 20 hours (c-c''), 7 days (d-d'') or 14 days (e-e'') of chase. (f) Bi-weekly incubations with 20nm Fluospheres for 15 days reveal the morphology of an endocytic cell in the gonad epithelium, schematized in (g). Magenta: Phalloidin (F-actin); blue: DAPI nuclear stain. DAPI+ vesicles are likely endocytic vesicles containing food-derived DNA. Mesenteries oriented with distal end to the top. Single confocal plane images of adult *Nematostella* mesenteries cross sections. Scale bars: (b-e)

25µm, (f) 10µm. CT: ciliated tract, CGT: cnidoglandular tract, IT: intermediate tract, ooc.: oocyte, put. mucus cell: putative mucus cell, SGE: somatic gonad epithelium, TT: trophic tract.

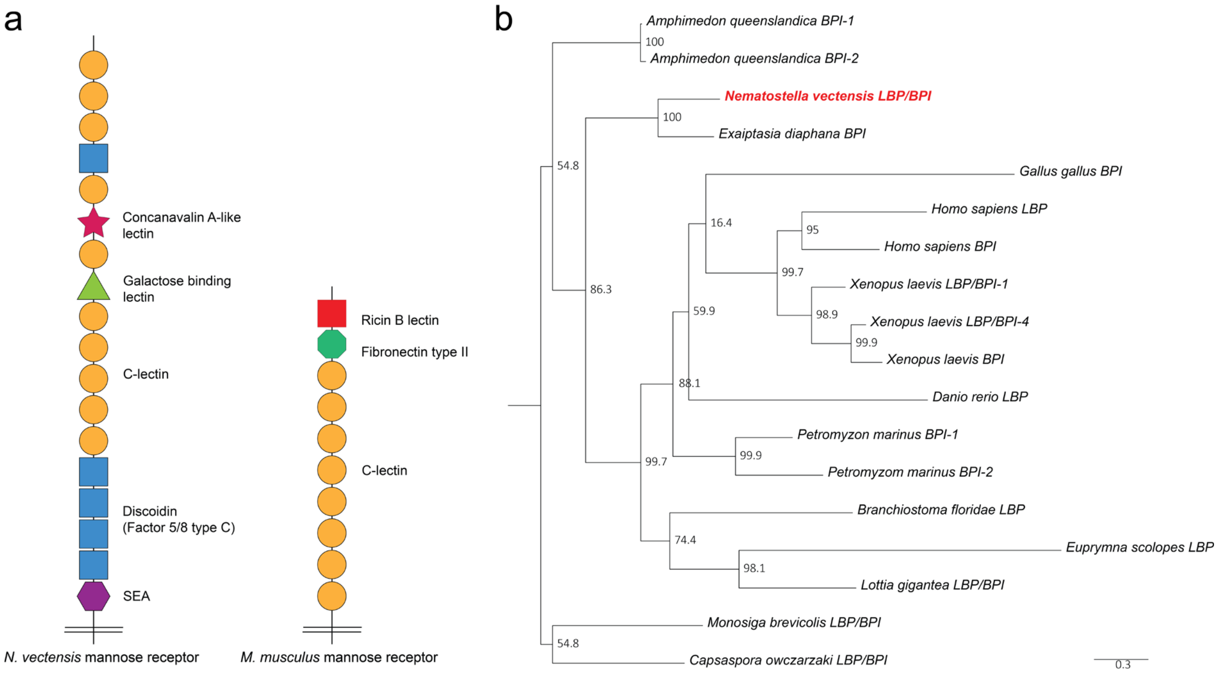


**Extended data figure 3**: ***N. vectensis*** **Mannose Receptor domain structure compared to its mammalian ortholog and phylogeny of the LBP/BPI gene orthologs.** (a) Predicted protein domains of the *N. vectensis* and *Mus musculus*. Orthology relies on sequence similarity between C-lectin domains. (b) Maximum likelihood tree (IQ-Tree) based on amino-acid alignments shows the orthology of the *N. vectensis* (highlighted in red) *LBP/BPI* gene. Node labels correspond to bootstrap values (1000 iterations). Scale indicates expected amino acid substitutions per site. *M. brevicolis* and *C. owczarzaki* were used as outgroups.

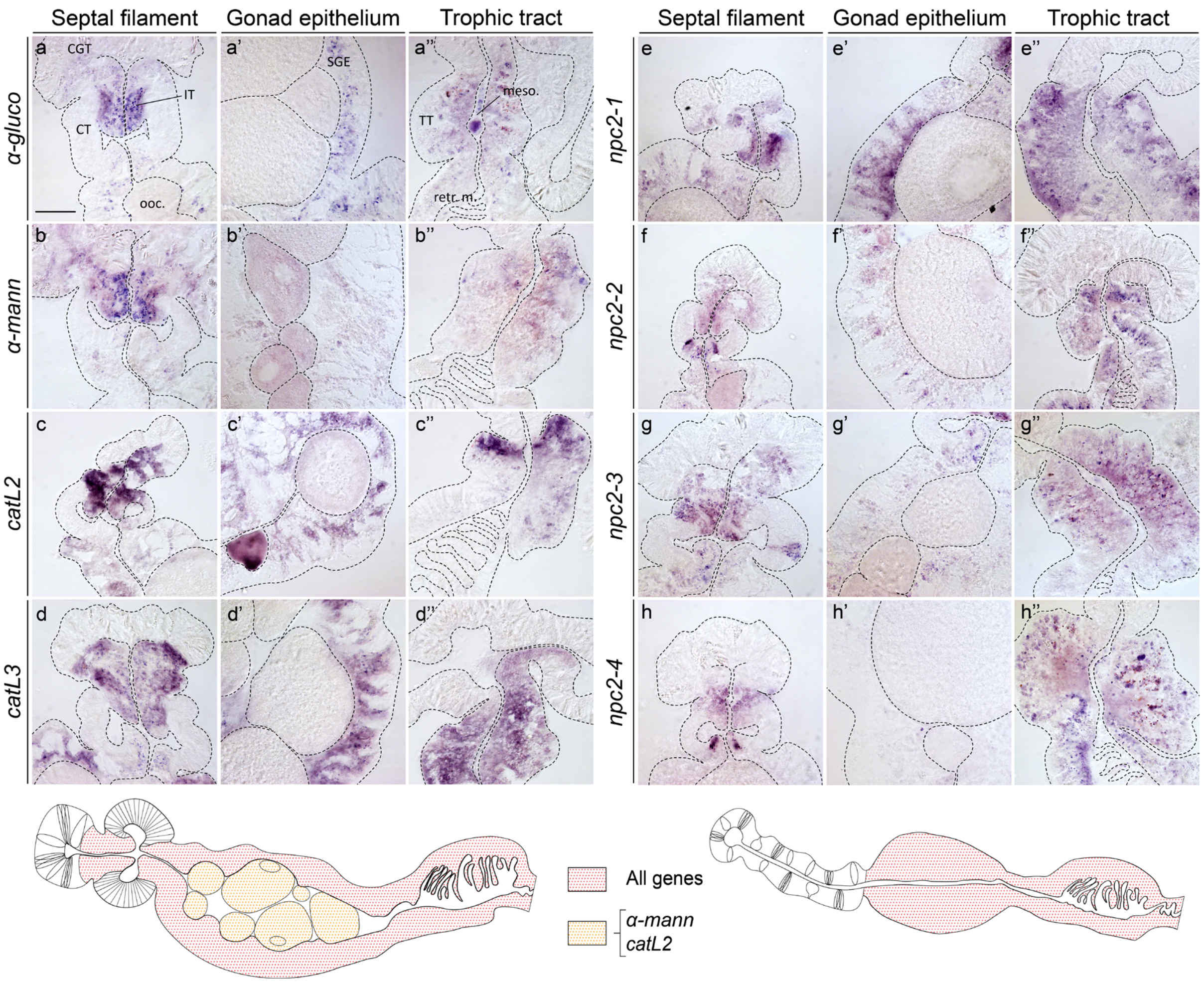


**Extended data figure 4: Expression of lysosomal enzyme and *npc2* cholesterol transporter genes in endocytic regions of the mesenteries.** mRNA expression of the lysosomal glycosidases *α-glucosidase (α-gluco)* (a-a'') and *α-mannosidase (α-mann)* (b-b''), the proteases *cathepsinL2* (*catL2)* (c-c'') and *cathepsinL3 (catL3)* (d-d'') and the cholesterol transporters *npc2-1* (e-e''), *npc2-2* (f-f''), *npc2-3* (g-g''), *npc2-4* (h-h'') in the septal filament (a-h), gonad epithelium (a'-h') and trophic tract (a''-h'') mesenterial regions. Cross sections of adult *N. vectensis* mesenteries stained by *in situ* hybridization. Mesenteries oriented with distal end to the top. Scale bar: 50µm. CT: ciliated tract, CGT: cnidoglandular tract, IT: intermediate tract, ooc.: oocyte, retr. m.: retractor muscle, RT: reticulate tract, SGE: somatic gonad epithelium, TT: trophic tract.

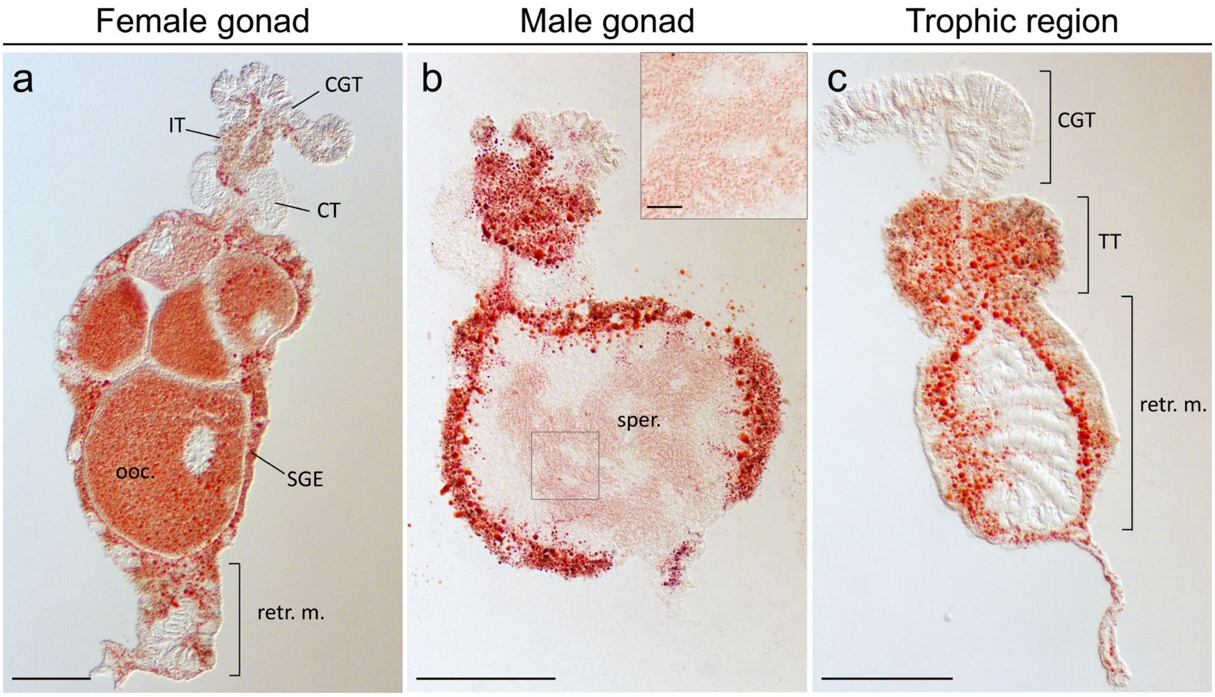


**Extended data figure 5: Abundant neutral lipid content in specific regions of the mesentery, including gametes.** (a-c) Oil Red O dye (ORO) shows abundant neutral lipid staining in the somatic gonad epithelium, retractor muscle and intermediate tract parts of the female (a) and male (b) gonad region, as well as in the trophic tract and retractor muscle of the trophic region (c). High lipid levels are found in oocytes (a). Sperm cells show distinct lipid vesicles (b, inset). Cross sections of adult Nematostella mesenteries after staining. All mesenteries oriented with distal end towards top. Scale bars: 100µm, except for close-up in (b): 10µm. CT: ciliated tract, CGT: cnidoglandular tract, IT: intermediate tract, ooc.: oocyte, retr. m.: retractor muscle, RT: reticulate tract, SGE: somatic gonad epithelium, sper.: spermary, TT: trophic tract.

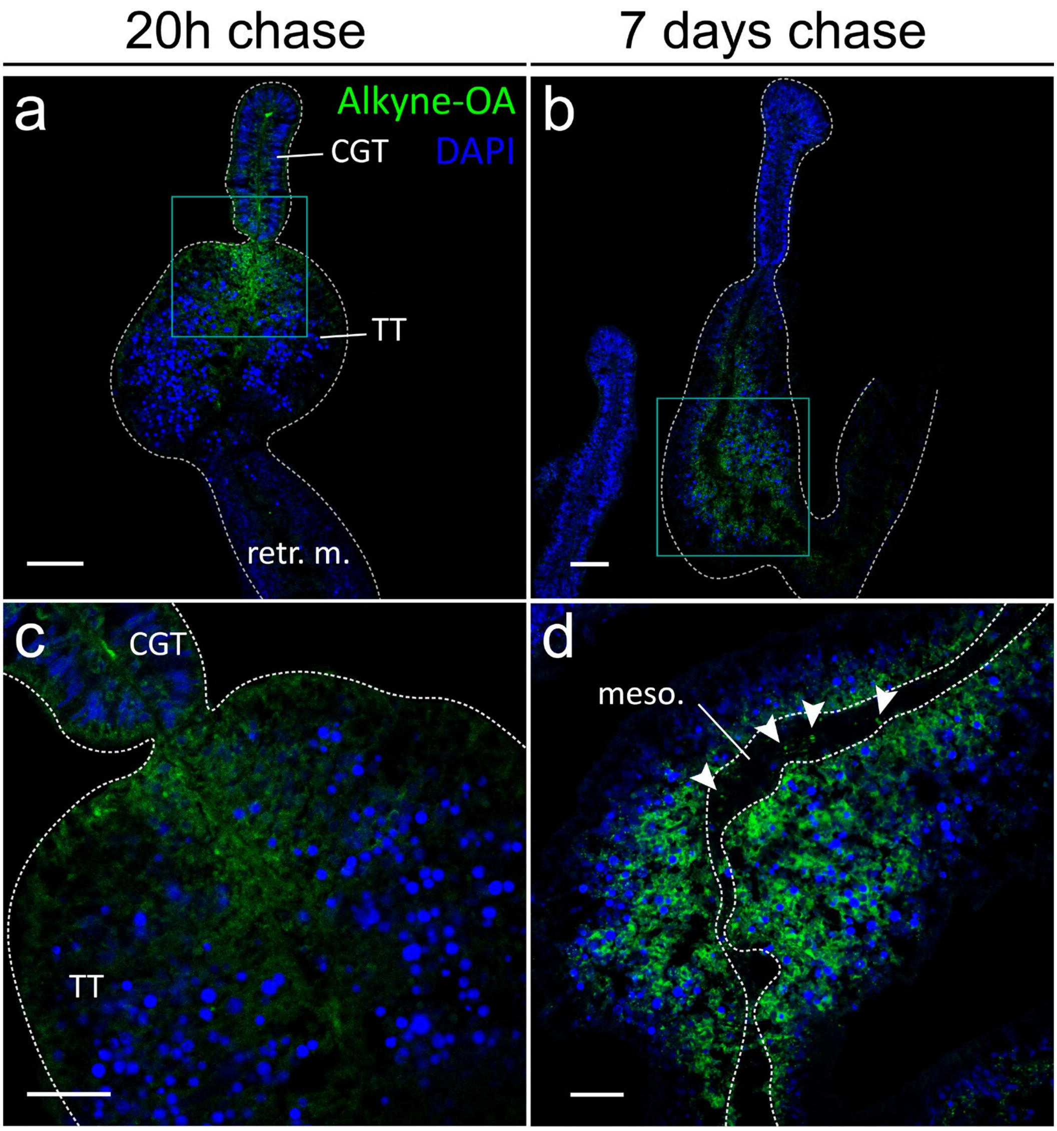


**Extended data figure 6: Uptake and translocation of alkyne-oleic acid (OA) in the trophic region.** Alkyne-OA localization in the trophic region after a 4-hours pulse of alkyne-oleic acid-enriched brine shrimps and a 20-hours (a, c) or 7-days chase period (b, d) (see Fig. 3a for experimental setup). Alkyne-OA (green) is present in the distal region of the trophic tract 20 hours post-ingestion of the enriched brine shrimps (a, boxed area enlarged in c), whereas after 7 days of chase, the signal is observed basally in cells of the central region of the trophic tract (b, boxed area enlarged in d) and within the mesoglea (d, white arrowheads). Scale bars: (a, b) 50µm, (c, d) 25µm. Single confocal plane images of adult female *N. vectensis* mesenteries cross sections. All mesenteries oriented with distal end towards top. Mesoglea is only demarcated when it could be unmistakably identified. CGT: cnidoglandular tract, meso.: mesoglea, retr. m.: retractor muscle, TT: trophic tract.

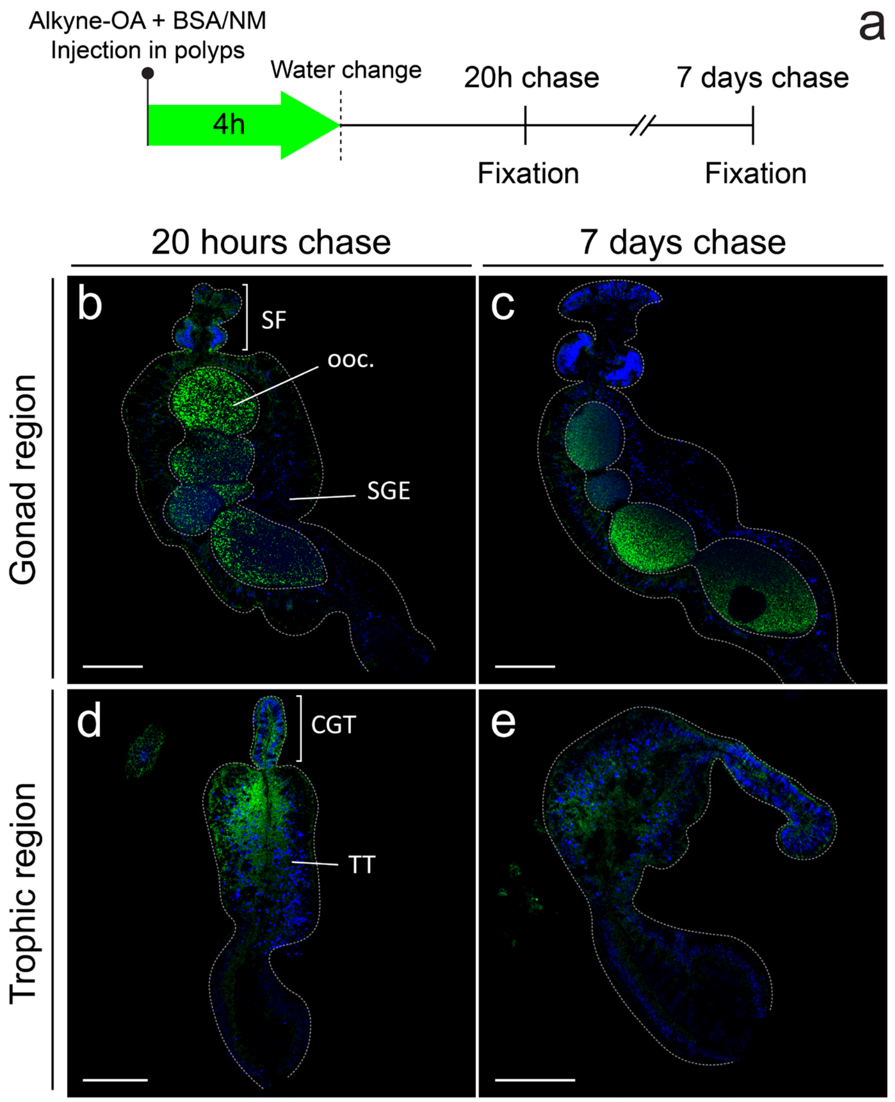


**Extended data figure 7: Alkyne-OA pulse-chase using BSA/NM suspension reproduces the results of alkyne-OA-enriched brine shrimp assay (Figure 1).** Alkyne-OA localization in the trophic region after a 4-hours pulse of BSA-coupled alkyne-OA and a 20-hours (b, d) or 7-days (c, e) chase (see a, experimental setup). Results are similar to those observed when using alkyne-OA-enriched brine shrimps (compare to Fig. 3b-e and Extended data fig. 5). Note for example similar staining of the distal trophic tract at 20 hours post-incubation (d), and the more basal location after 7 days of chase (e). In the gonad region, stained vesicles are present in the gonad epithelium after 20 hours (b) but absent after 7 days (c). Alkyne-OA is present in the oocytes in both conditions (b, c). Single confocal plane images of adult female *Nematostella* mesenteries cross sections. Mesenteries oriented with distal end towards top. Scale bars: 100µm. CGT: cnidoglandular tract, ooc.: oocyte, SGE: somatic gonad epithelium, SF: septal filament, TT: trophic tract.

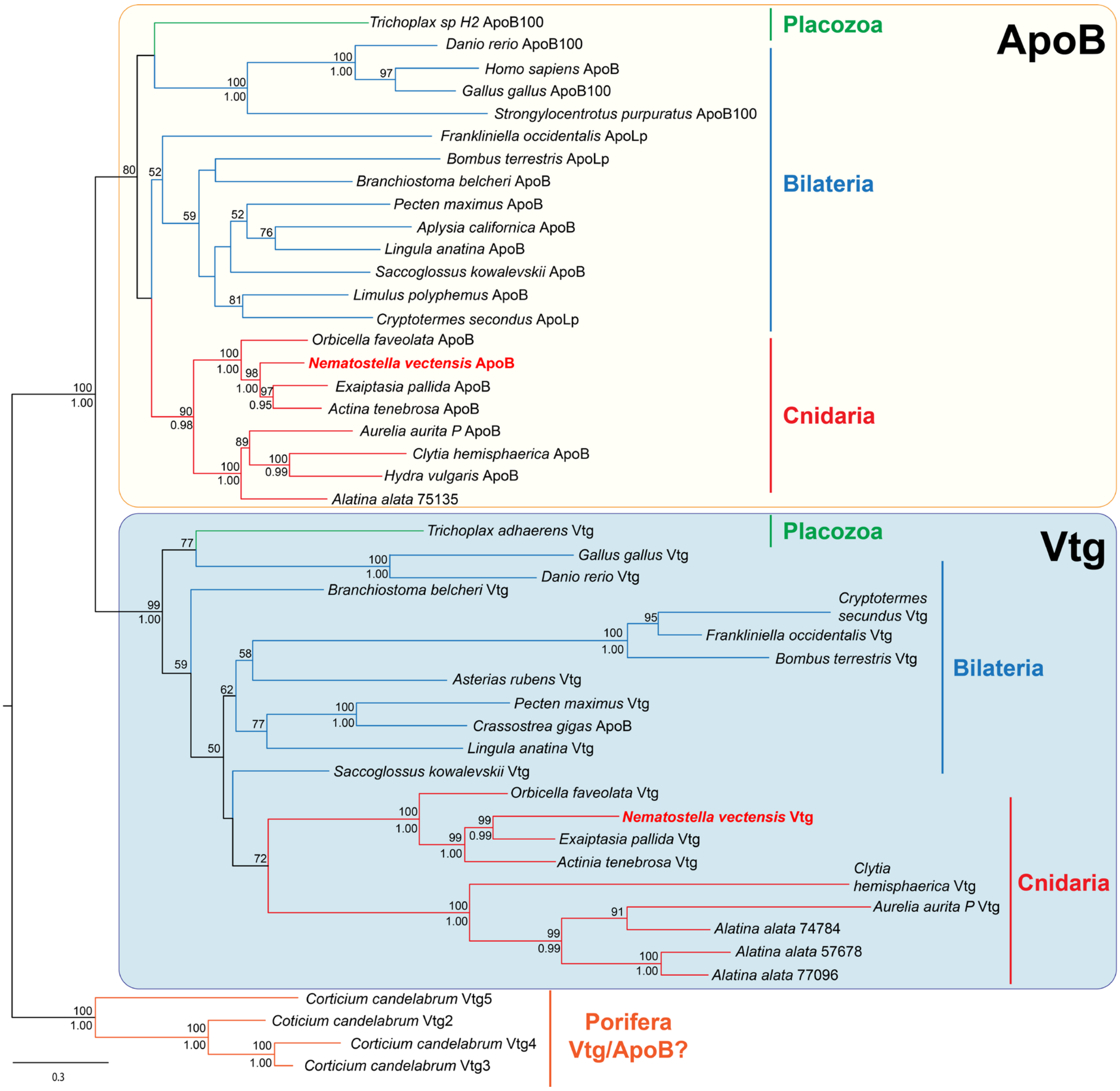


**Extended data Figure 8:** ***Nematostella*** **possesses** ***single apoB*** **and** ***vtg*** **gene orthologs.** Maximum likelihood tree (IQ-Tree) based on amino acid alignments shows the orthology of the *N. vectensis* (highlighted in red) ApoB and Vtg with corresponding bilaterian, cnidarians and placozoan proteins. Node labels correspond to bootstrap percentages (above branch) or Bayesian probabilities (below branch). Scale indicates amino acid substitutions per site. Note that the orthology of sponge proteins to Vtg or ApoB remains undetermined. ApoLp: apolipophorin.

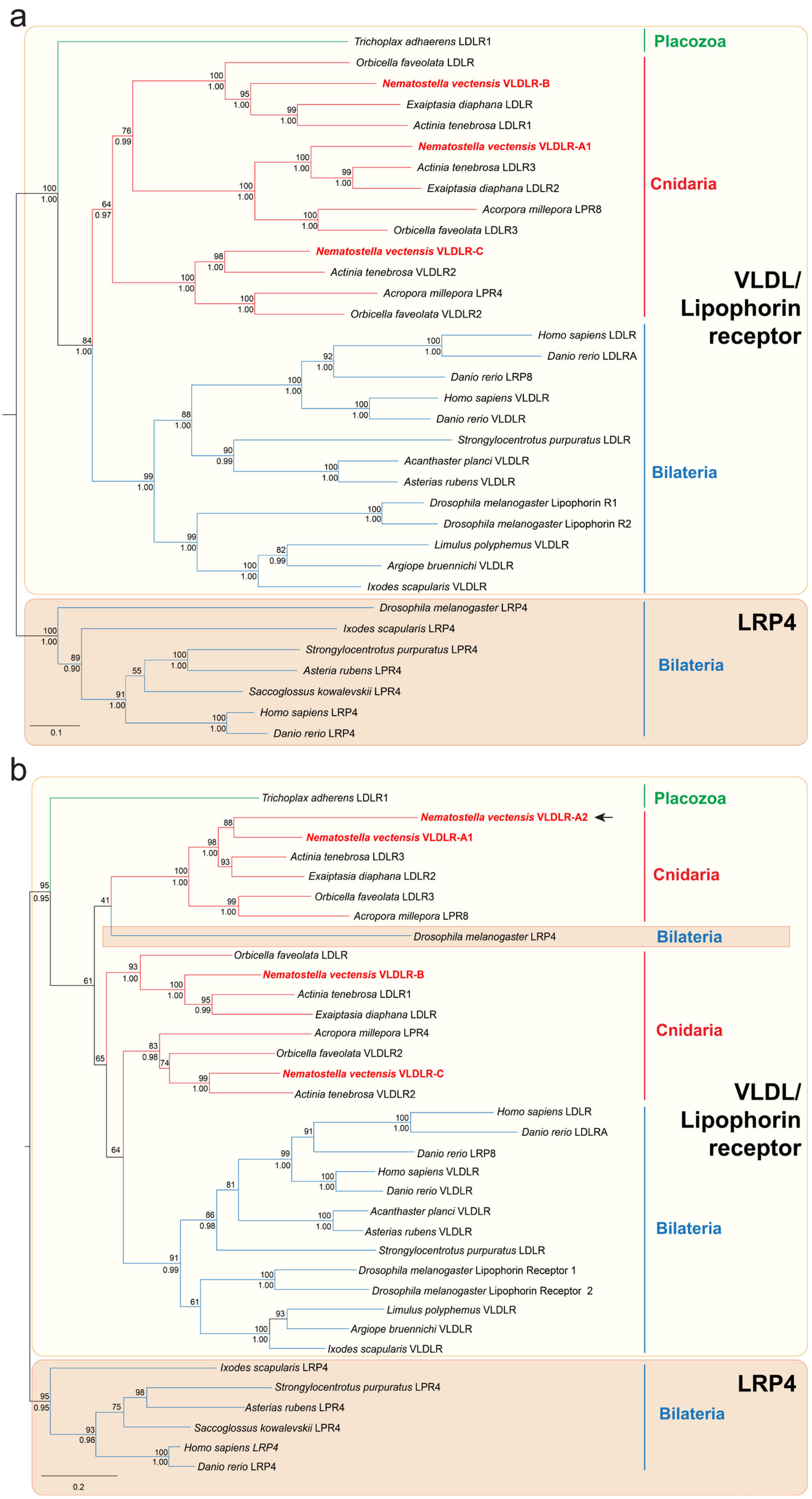
a
Trichoplax adhaerens LDLR1
Orbicella faveolata LDLR
Nematostella vectensis VLDLR-B
Exaiptasia diaphana LDLR
Actinia tenebrosa LDLR1
Nematostella vectensis VLDLR-A1
Actinia tenebrosa LDLR3
Exaiptasia diaphana LDLR2
Acorpora millepora LPR8
Orbicella faveolata LDLR3
Nematostella vectensis VLDLR-C
Actinia tenebrosa VLDLR2
Acropora millepora LPR4
Orbicella faveolata VLDLR2
Homo sapiens LDLR
Danio rerio LDLRA
Danio rerio LRP8
Homo sapiens VLDLR
Danio rerio VLDLR
Strongylocentrotus purpuratus LDLR
Acanthaster planci VLDLR
Asterias rubens VLDLR
Drosophila melanogaster Lipophorin R1
Drosophila melanogaster Lipophorin R2
Limulus polyphemus VLDLR
Argiope bruennichi VLDLR
Ixodes scapularis VLDLR
Drosophila melanogaster LRP4
Ixodes scapularis LRP4
Strongylocentrotus purpuratus LPR4
Asteria rubens LPR4
Saccoglossus kowalevskii LPR4
Homo sapiens LRP4
Danio rerio LRP4
0.1
Placozoa
Cnidaria
VLDL/
Lipophorin
receptor
Bilateria
LRP4
Bilateria
b
Trichoplax adherens LDLR1
Nematostella vectensis VLDLR-A2
Nematostella vectensis VLDLR-A1
Actinia tenebrosa LDLR3
Exaiptasia diaphana LDLR2
Orbicella faveolata LDLR3
Acropora millepora LPR8
Drosophila melanogaster LRP4
Orbicella faveolata LDLR
Nematostella vectensis VLDLR-B
Actinia tenebrosa LDLR1
Exaiptasia diaphana LDLR
Acropora millepora LPR4
Orbicella faveolata VLDLR2
Nematostella vectensis VLDLR-C
Actinia tenebrosa VLDLR2
Homo sapiens LDLR
Danio rerio LDLRA
Danio rerio LRP8
Homo sapiens VLDLR
Danio rerio VLDLR
Acanthaster planci VLDLR
Asterias rubens VLDLR
Strongylocentrotus purpuratus LDLR
Drosophila melanogaster Lipophorin Receptor 1
Drosophila melanogaster Lipophorin Receptor 2
Limulus polyphemus VLDLR
Argiope bruennichi VLDLR
Ixodes scapularis VLDLR
Ixodes scapularis LRP4
Strongylocentrotus purpuratus LPR4
Asterias rubens LPR4
Saccoglossus kowalevskii LPR4
Homo sapiens LRP4
Danio rerio LPR4
0.2
Placozoa
Cnidaria
Bilateria
Cnidaria
VLDL/
Lipophorin
receptor
Bilateria
LRP4
Bilateria

**Extended data figure 9: *Nematostella* has four VLDL/Lipophorin receptor protein paralogs with VLDLR-A1, -B and -C having orthologs in other anthozoans (a) and VLDLR-A2 being a truncated, *Nematostella*-specific paralog of VLDLR-A1 (b).** Maximum likelihood trees (IQ-Tree) based on amino acid alignments shows the orthology of four Nematostella (highlighted in red) Very Low-Density Lipoprotein Receptors (VLDLRs) with the corresponding bilaterian proteins. +LRP4 proteins, which are the closest-related LDLR subfamily, are used as an outgroup. Tree in (b) is based on a short, manually curated amino acid alignment including only the positions aligning with the truncated Nv-VLDLR-A2 protein sequence and shows the clear orthology of VLDLR-A2 as a paralog of the Nv-VLDLR-A1 protein. Node labels correspond to bootstrap percentages (above branch) or Bayesian probabilities (below branch). Scale indicates expected amino acid substitutions per site. LRP4 proteins, which are the closest-related LDLR subfamily, are used as an outgroup. Lipophorin R: lipophorins receptor.
